# A multiadjuvant polysaccharide-amino acid-lipid (PAL) subunit nanovaccine generates robust systemic and lung-specific mucosal immune responses against SARS-CoV-2 in mice

**DOI:** 10.1101/2023.05.05.539395

**Authors:** Bhawana Pandey, Zhengying Wang, Angela Jimenez, Eshant Bhatia, Ritika Jain, Alexander Beach, Drishti Maniar, Justin Hosten, Laura O’Farrell, Casey Vantucci, David Hur, Richard Noel, Rachel Ringuist, Clinton Smith, Miguel A. Ochoa, Krishnendu Roy

## Abstract

Existing parenteral SARS-CoV-2 vaccines produce only limited mucosal responses, which are essential for reducing transmission and achieving sterilizing immunity. Appropriately designed mucosal boosters could overcome the shortcomings of parenteral vaccines and enhance pre-existing systemic immunity. Here we present a new protein subunit nanovaccine using multiadjuvanted (e.g. RIG-I: PUUC, TLR9: CpG) polysaccharide-amino acid-lipid nanoparticles (PAL-NPs) that can be delivered both intramuscularly (IM) and intranasally (IN) to generate balanced mucosal-systemic SARS-CoV-2 immunity. Mice receiving IM-Prime PUUC+CpG PAL- NPs, followed by an IN-Boost, developed high levels of IgA, IgG, and cellular immunity in the lung, and showed robust systemic humoral immunity. Interestingly, as a purely intranasal vaccine (IN-Prime/IN-Boost), PUUC+CpG PAL-NPs induced stronger lung-specific T cell immunity than IM-Prime/IN-Boost, and a comparable IgA and neutralizing antibodies, although with a lower systemic antibody response, indicating that a fully mucosal delivery route for SARS-CoV-2 vaccination may also be feasible. Our data suggest that PUUC+CpG PAL-NP subunit vaccine is a promising candidate for generating SARS-CoV-2 specific mucosal immunity.

## INTRODUCTION

The COVID-19 pandemic continues to cause a global health crisis with frequent viral mutations and uncontrolled transmission in many parts of the world. In the U.S. alone, weekly death rates from COVID-19 remain well over 1000, even three years after the start of the pandemic (*1*). Almost 72% of the world’s population have received the licensed parenteral mRNA-LNP and adenoviral vectors-based vaccines (*1–3*). These approved vaccines are designed to be administered via intramuscular (IM) route, show high efficacy against severe SARS-CoV-2 infections, and reduce hospitalization and deaths (*2–4*). Unfortunately, recent studies have shown that even after vaccination with booster doses, there are asymptomatic, symptomatic, and some severe cases of SARS-CoV-2 infection observed after a few months. These results indicate declining antiviral immunity within a short period and raise questions about durable efficacy and protection through these vaccines (*5, 6*). Moreover, continuous viral mutation (VOC: variants of concern) that evades the immune response also results in reduced vaccine effectiveness (*7, 8*). Additionally, vaccine accessibility and acceptance remain a significant concern (*6*).

The SARS-CoV-2 pathogen infects human cells through binding of its RBD (Receptor Binding Domain) region to the ACE-2 receptors present in the cells of respiratory tract (mucosal tissues), which makes SARS-CoV-2 primarily a mucosal pathogen (*9*). However, IM vaccinations predominantly induce systemic immune responses (circulating antibodies, memory B cells, effector T cells), with limited mucosal immunity at the sites of infection, i.e., nasopharynx and lungs (*10, 11*). The immune response generated after IM immunization leaves the upper respiratory tract vulnerable to viral replication and dissemination, leading to reduced sterilizing immunity through IM vaccines (*12, 13*). Collectively, approved vaccines have limited efficacy in preventing SARS-CoV-2 infection due to several factors, including poor respiratory mucosal immunity, viral immune evasion, and increased viral transmission. Mucosal vaccination could be one of the potential solutions to the above problems. It may generate both mucosal and systemic antiviral immune responses (humoral and cellular) similar to a natural infection and may ultimately lead to better protection and reduced transmission (*14, 15*).

The antiviral mucosal immunity is characterized by the generation of robust mucosal IgA, IgG, and neutralizing antibodies in the bronchoalveolar lavage (BAL) fluid, lung-specific T cell immunity (Tissue-resident memory T and B cell) with TH1 type and systemic immune responses (*14–17*). Vaccine administration through the respiratory route (nasal/oral vaccines) generates more potent tissue-level mucosal immunity against SARS-CoV-2, compared to the IM route, thus emerged as an effective approach for new mucosal vaccines (*18, 19*). One can either boost the already IM vaccinated population with the intranasal vaccine (heterologous vaccine strategy), which may also strengthen the circulating immunity achieved via intramuscular priming (IM-Prime/IN-Boost), or use them for both priming and boosting (homologous strategy, IN-Prime/IN-Boost) in the unvaccinated population, which could increase compliance and acceptance (*18–23*). Recently, encouraging preclinical findings have emerged utilizing the IM-Prime/IN-Boost approach. One study involved systemic priming with an mRNA-LNP vaccine followed by an intranasal boost of spike protein, while another study administered a Diphtheria toxin-conjugated vaccine through IM-Prime and IN-Boost (*21, 22*). This vaccination strategy may enhance systemic immunity through mucosal boost (a prime-pull mechanism) and help achieve sterilizing immunity against SARS-CoV-2.

As of December 2022, among a total of 21 mucosal vaccines in trials, five have been authorized for use or registered for regulatory agency review for SARS-CoV-2. Out of which, three are viral vector-based vaccines: Bharat Biotech in India, Gamaeleya in Russia, and CanSino Biologics in China (*24, 25*). However, the effectiveness of these viral vector-based vaccines for the worldwide population is still under assessment. Indeed, the fact that none of these vaccines have been authorized for use in the United States or Europe, nor has the WHO granted them emergency use listing, could be attributed to the limited data available from clinical efficacy trials to assess their impact on transmission, infection, or disease (*26*). In rare cases, for live-attenuated vaccines, there might be a potential risk of returning to their virulent state (*26*). Mucosal vaccines for influenza that utilized intranasal live attenuated pathogens had been approved previously but were not 100% effective and safe (*27*). Therefore, there is a need to develop new mucosal vaccine strategies that not only overcome the limitations of IM vaccines and achieve high levels of durable protection against infection but also mitigate the post-COVID severe effects, such as post-acute sequelae of SARS-CoV-2 infection and long COVID.

Protein subunit vaccines are known to be comparatively safer as they use only a fragment of a pathogen. Some promising results with recombinant protein subunit vaccines have been recently reported (*28, 29*). However, protein subunit vaccines exhibit less immunogenicity (*30*) requiring the use of vaccine adjuvants, which are often delivered using biomaterial-based polymeric nanoparticle (NP) formulations, often known as nanovaccines (*31, 32*). Adjuvants on nanoparticles increase antigen immunogenicity by activating pattern recognition receptors (PRRs) of innate immune system, modulating the antigen pharmacokinetics, and facilitating antigen dose sparing (*33, 34*). Several studies have shown that combination of adjuvants in a vaccine formulation can elicit synergistic, complementary, and antagonistic effects on innate and adaptive immunity (*34*). Several adjuvants that target multiple PRRs, including RLRs (retinoic inducible gene 1: (RIG-I)-like receptors) and Toll-like receptors (TLRs: 4, 7/8, 9), either alone or in combination, have been utilized in studies related to SARS-CoV-2 mucosal vaccines. While these adjuvants have shown promising results in animal models, they are not able to provide complete protection against infection (*13, 21, 35–37*). However, RIG-I agonists are mainly used to enhance antiviral immunity in other viral infections such as influenza or the west nile virus (*38–41*).

Here we designed a novel, multiadjuvanated protein subunit SARS-CoV-2 nanovaccine formulation using different combinations of PRR-agonists-PUUC RNA: RLRs agonist, CpG DNA: TLR9 agonist and R848: TLR7/8 agonist, which are loaded on degradable, polysaccharide- amino acid-lipid polymeric nanoparticles (PAL-NPs). We first evaluated the best combination adjuvant formulation that would potentially enhance both the mucosal and systemic immunity against SARS-CoV-2 via the intramuscular (IM) prime and intranasal (IN) boost (IM-Prime/IN-Boost) strategy. We found that PAL-NPs with PUUC (RIG-I agonist) and CpG DNA (TLR9 agonist), combined with the recombinant stabilized Spike S1 trimer protein, induced robust mucosal and systemic humoral and local cellular immunity (TH1 response) with IM-Prime/IN-Boost group. However, the IN-Prime/IN-Boost group also induces robust lung T cell-mediated immunity, higher than the IM-Prime/IN-Boost group, and a comparable mucosal humoral response (IgA and neutralizing antibodies), which suggests that a mucosal delivery route could be attainable for future vaccines compared to only parenteral route.

## RESULTS

### Synthesis and characterization of multiadjuvanated PAL-NP vaccine formulations

We developed a multiadjuvanted nanoparticle-based SARS-CoV-2 protein subunit vaccine using a newly designed polymer-lipid molecule (**Fig. 1**). We have previously reported the use of polymer particles for delivery of combination adjuvants and antigens, both in vitro and in vivo (*38, 42, 43*). Proper selection of polymers is necessary for better loading and delivery of multiple charged or hydrophobic adjuvants on the polymer nanoparticles. Synthetic amphiphilic polymers can fulfill these criteria because they are chemically modified with the desired functional group moieties, specific for adjuvant loading and delivery. Therefore, we first synthesized the amphiphilic polymer-lipid [OCMC-S-S-(A/H)-SA], namely- polysaccharide-amino acid-lipid (PAL polymer), by sequential, structural, and chemical modification of polysaccharide (chitosan Mw: 15 KDa) (**Fig. 1A, fig. S1A, and supplementary text 2.1**). The PAL polymer synthetic steps start with (i) carboxylation at C-6 position using mono-chloroacetic acid in slight basic medium, (ii) thiolation of C-2 amine groups with thioglycolic acid using carbodiimide chemistry, (iii) disulfide formation between thiols at C-2 position of carboxylated chitosan and the thiols of cysteamine, (iv) carbodiimide conjugation of the carboxyl group of N-α Boc protected amino acids (arginine and histidine) with amine group of cysteamine (v) stearyl amine conjugation at C-6 carboxyl group (O-substitution) using carbodiimide chemistry, (vi) final deprotection of Boc groups. Purification of all steps was performed by dialysis or precipitation and followed by lyophilization at the required pH. The incorporation of functional groups in polysaccharides is confirmed by proton NMR spectroscopy. We confirm the conjugation and incorporation of amino acids and stearyl chains (lipid) into the chitosan backbone by comparing the ^1^HNMR spectra of thiolated chitosan polysaccharide and the final modified polysaccharide-amino acid-lipid polymer (**Fig. 1B**). The appearance of a characteristic peak at 7.1-7.4 ppm is due to a guanidine functional group in arginine, suggesting the successful arginine grafting in polysaccharide. Additionally, a characteristic peak of imidazole ring protons at 8.6 ppm and 7.6 ppm confirms histidine grafting on the polysaccharide backbone. The peaks at 1.2 ppm (-CH2-) and 1.6 ppm (-CH3) in the ^1^H NMR spectrum of amphiphilic chitosan polymer confirm the successful incorporation of the stearyl chain in chitosan. The presence of disulfide bond formation was confirmed by Elmann’s assay, which shows a total reduction in free thiol concentration after cysteamine conjugation (**fig. S1B**).

**Fig. 1.**
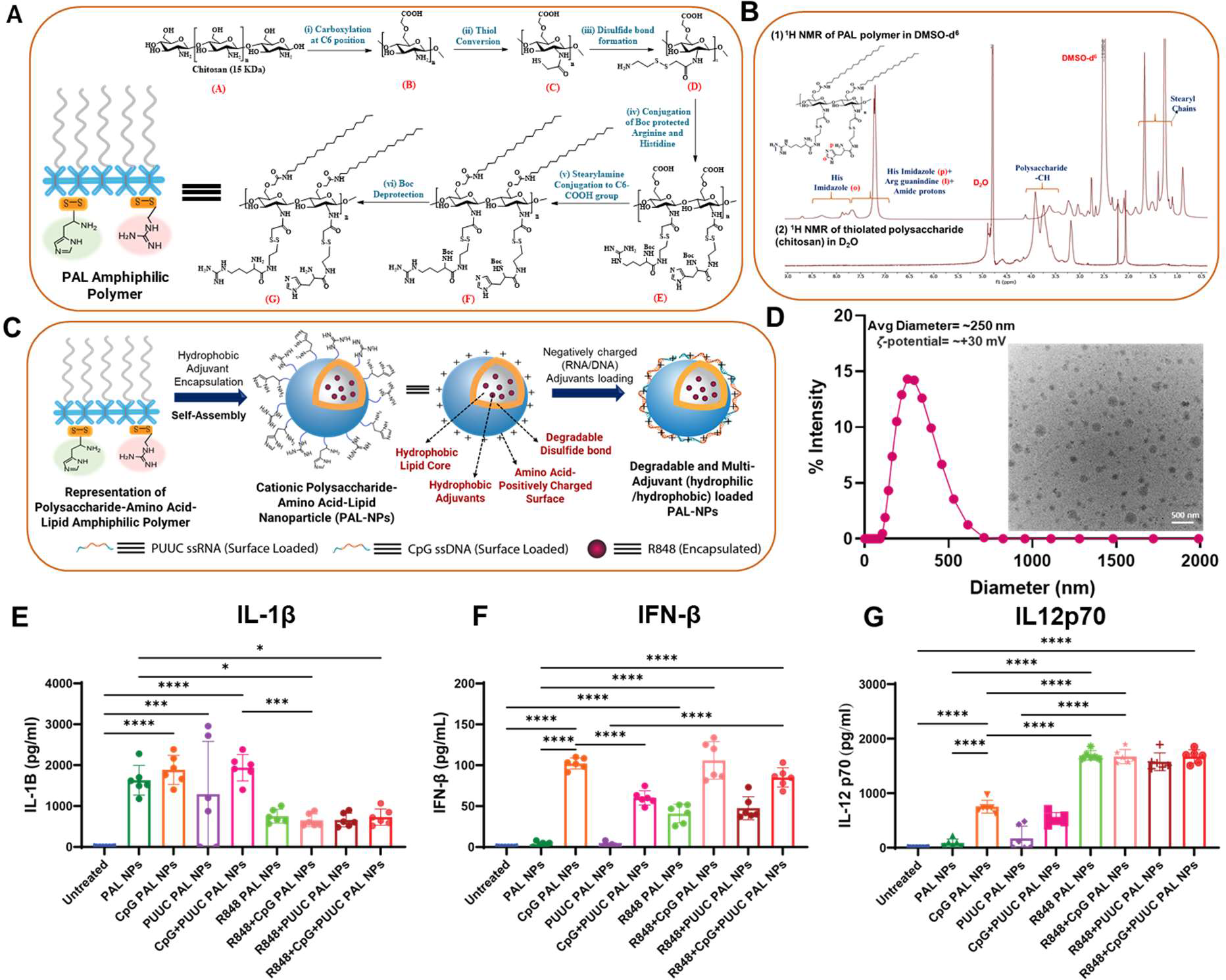
Synthesis and characterization of multiadjuvanated PAL-NPs. (**A**) Multistep synthetic scheme of cationic and degradable polysaccharide-amino acid-lipid (PAL) amphiphilic polymer. (**B**) Comparison of ^1^H NMR spectra of amphiphilic polymer with the ^1^H NMR spectra of thiolated chitosan polymer after structural modification. (**C**) Schematic of PAL-NPs fabrication from polymer, depiction of PAL-NPs with encapsulated hydrophobic adjuvant (R848) and surface- loaded nucleic acids adjuvant (PUUC, CpG) for their delivery in both in vitro and in vivo. (**D**) Physiochemical characterization of PAL-NPs: hydrodynamic diameter and zeta potential, (inset: TEM image of PAL-NPs, scale bar is 500 nm). (**E** to **F**) Nanoparticle co-delivery of multiadjuvants broadens the innate immune response in GM-CSF differentiated murine BMDCs. Murine GM- CSF differentiated BMDCs were treated with single/dual/triple adjuvanated PAL-NP formulations and controls. Analysis of cytokine level: IL-1β (**E**), IFN-β (**F**), and IL12p70 (**G**) after 24 h of adjuvanted PAL-NP treatment (*n* = 6) from GM-CSF differentiated murine BMDCs. Error bars represent SEM (standard error of the mean). Statistical significance was determined by one-way ANOVA followed by Tukey’s post-hoc test for multiple comparisons. *p ≤ 0.05, **p ≤ 0.01, ***p ≤ 0.001, ****p ≤ 0.0001 for all graphs.

We further synthesized a cationic and degradable nanoparticle from amphiphilic PAL polymer, known as PAL-NP, using the probe sonication method in DMSO/water mixture and purification by dialysis. Multiadjuvanated PAL-NPs were prepared by the loading of different adjuvant combinations of PUUC RNA (targeting RIG-I-like receptors: retinoic acid-inducible gene I), CpG DNA (targeting TLR9) and R848 (targeting TLR7/8), either by surface electrostatic adsorption (anionic adjuvants: PUUC and CpG) or by encapsulation (hydrophobic adjuvants: R848) inside the nanoparticles’ lipid core (**Fig. 1C and table S1**). The blank and R848 loaded PAL-NPs have an average hydrodynamic size of ∼250 nm and zeta potential of ∼ +30 mV at pH ∼7.0, which makes it appropriate for surface loading of nucleic acid adjuvants (**Fig. 1D**). Transmission electron microscopy (TEM) was used to determine the morphology of PAL-NPs, revealing a well-dispersed spherical shape with the average diameter between 150-200 nm (**Fig. 1D, inset**). The degradability of PAL-NPs mediated by disulfide bond reduction was confirmed by a decrease in nanoparticle size from ∼200 nm to ∼50 nm with the addition of dithiothreitol (DTT) for 24 h (**fig. S1C**), which was not observed in disulfide-bond deficient PAL-NPs (non-degradable PAL-NPs).

To examine the immunostimulatory effects of different combinations of PRR agonists (RIG-I agonist and TLRs agonist) in vitro, we screened eight PAL-NPs adjuvant formulations (CpG, PUUC, CpG+PUUC, R848, R848+CpG, R848+PUUC, R848+CpG+PUUC) (**Fig. 1, E to G**). These formulations were incubated with murine bone-marrow-derived dendritic cells (BMDCs) generated using GM-CSF cytokines for 7 days in a 96-well plate (**table S1**). After 24 h, the collected supernatants were analyzed to quantify proinflammatory cytokine secretion. Preliminary in vitro studies show that PUUC+CpG PAL-NPs significantly increased the secretion of proinflammatory cytokine IL-1β (**Fig. 1E**). However, IL-1β secretion is mainly driven by CpG PAL-NPs. Similarly, the IFN-β secretion is also CpG driven, but interestingly, R848+CpG PAL- NPs synergistically increase the IFN-β secretion (**Fig. 1F**). R848 PAL-NPs is the only group that significantly stimulates the secretion of proinflammatory cytokine IL12p70 (**Fig. 1G**). Furthermore, none of the triple adjuvanated PAL-NPs enhanced proinflammatory cytokine secretion. Surprisingly, only blank PAL-NP, which are considered as a control group due to the absence of real RLR and TLR agonist adjuvants, also show considerable IL-1β secretion but do not stimulate the secretion of the IFN-β and IL12p70. However, both PUUC+CpG PAL-NPs and R848+CpG PAL-NPs enhance the stimulation of the initial innate immune response. These results inspired us to evaluate whether these RIG-I and TLR-targeted PAL-NPs would enhance the immune responses for SARS-CoV-2 in vivo.

### PAL-NPs protein subunit vaccine adjuvanated with RIG-I (PUUC) and TLR9 (CpG) agonists elicit robust SARS-CoV-2 mucosal and systemic humoral immune responses, when delivered IM-Prime/IN-Boost

For boosting the current SARS-CoV-2 vaccine or developing a mucosal vaccine, an ideal vaccine candidate should have an appropriate adjuvant combination that generates potent and balanced mucosal and systemic immunity. Therefore, we first performed the in vivo screening of multiple adjuvant combinations on PAL-NPs and evaluated the best adjuvant combination that enhances the mucosal and systemic SARS-CoV-2 immune response. We selected four adjuvanated (PUUC, R848, R848+CpG, PUUC+CpG) PAL-NPs groups and administered them in mice through IM-Prime/IN-Boost strategy (**Fig. 2A**). These nanovaccine formulations were prepared by loading/encapsulation the adjuvants (PUUC, R848, R848+CpG, PUUC+CpG) on PAL-NPs and mixing them with stabilized recombinant SARS-CoV-2 S1 trimer subunit as the target antigen. S1 trimer subunit is more immunogenic than the RBD alone due to the presence of other epitopes at the outer part of the RBD, which contribute to the neutralization (*44*). The blank NPs (no real adjuvants) in our study exhibit minimal immune responses; therefore, we have considered them as the control group along with PBS. Mice were immunized via the IM-Prime at day 0 and boosted via IN route at day 21 with PAL-NPs vaccine formulations and sacrificed at day 35 (14 days post-boost), and BAL fluid and serum samples were collected for analyzing the generated mucosal and systemic humoral response. We first quantified the mucosal anti-spike S1 IgA and IgG levels in the BAL fluid. We found that PUUC+CpG PAL-NP group significantly increases BAL anti-spike IgA and IgG levels at 1:10 dilution compared to other formulations, including control groups (**Fig. 2, B and C**). The secreted mucosal IgA and IgG levels in BAL fluid of mice vaccinated with adjuvanated PAL-NPs vaccine formulations and controls follow the order: PUUC+CpG>PUUC>R848>R848+CpG>PAL-NPs>PBS. To assess TH1/TH2 skewed IgG response in BAL fluid elicited through adjuvanated PAL-NPs vaccine immunization, we evaluated IgG subsets (IgG1 and IgG2a) present (**Fig. 2, D and E**). The TH1-associated IgG2a levels were significantly higher in mice immunized with PUUC+CpG PAL-NP group than in the other groups (**Fig. 4E**). The IgG2a/IgG1 ratio suggests that both PUUC+CpG and R848+CpG groups show the TH1-biased response, but a comparatively higher response was observed in the former (**Fig. 2F**). This data indicates that PUUC+CpG PAL-NPs vaccine formulation generates more potent mucosal IgA and IgG in BAL fluid.

**Fig. 2.**
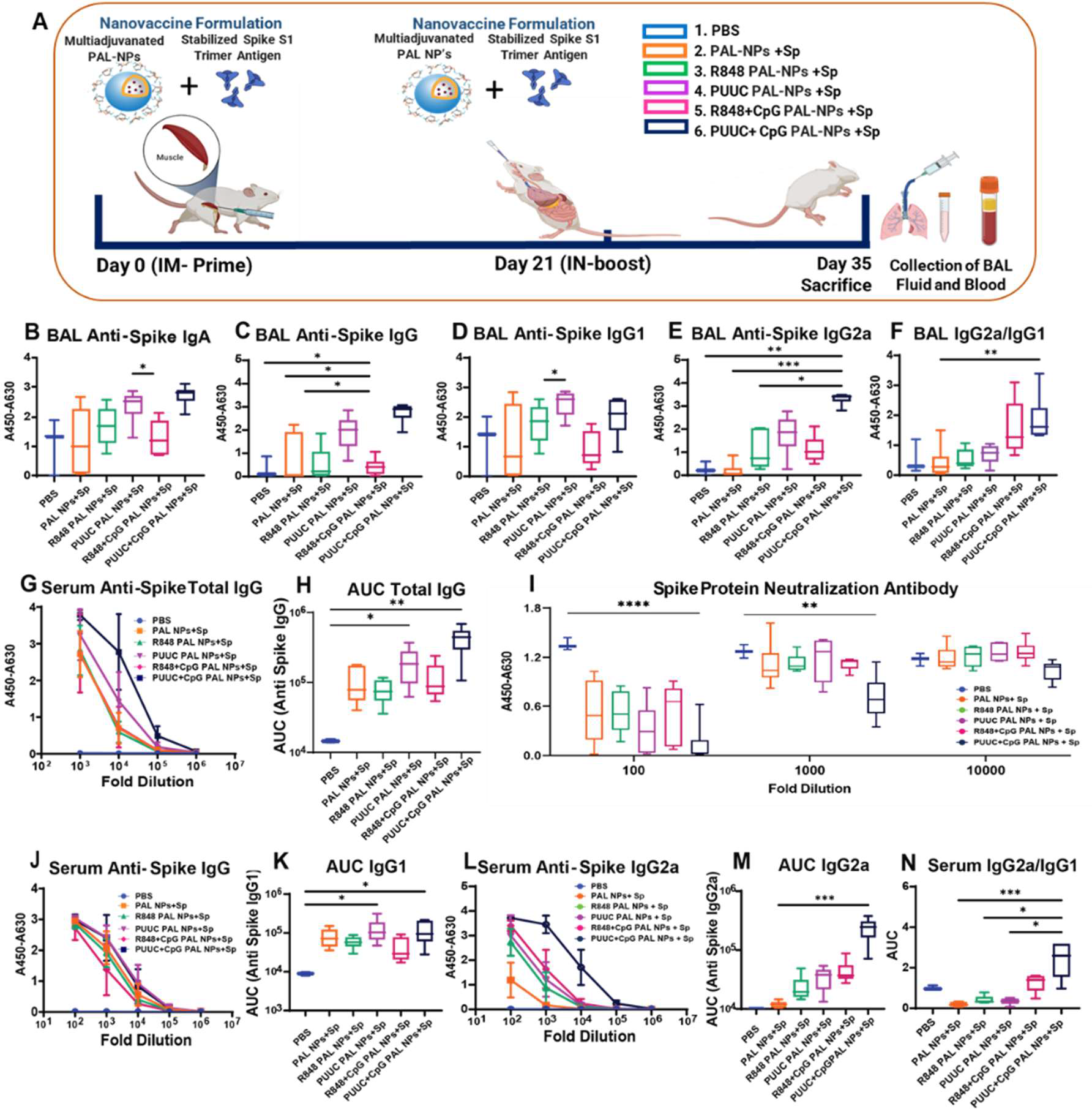
A subunit nanovaccine formulation of PAL-NPs adjuvanated with RIG-I (PUUC) and TLR9 (CpG) agonists mixed with S1 spike protein elicits robust SARS-CoV-2 mucosal and systemic humoral immunity, when delivered IM-Prime/IN-Boost. (**A**) Experimental schematics: Female BALB/c mice (n=3 for PBS and n=6 for other adjuvanated PAL-NP formulations) were immunized IM into both anterior tibialis muscles at day 0 (1^st^ dose) with vaccine formulation of adjuvanated PAL-NPs (NPs: 250 µg, PUUC: 20 µg, CpG: 40 µg and R848: 20 µg) and combined with stabilized spike (Sp) S1 trimer protein at a dose of 1000 ng respectively. On day 21, mice received the 2^nd^ dose of vaccine formulation IN using similar doses of adjuvants, PAL-NPs, and spike protein, except for the CpG dose reduced to 20 µg. Mice were euthanized after two weeks on day 35 to collect BAL fluid and serum. BAL and serum from vaccinated mice were assayed with ELISA assay. (**B** to **E**) BAL fluid from vaccinated mice was assayed for anti- spike IgA (**B**), IgG (**C**), IgG (**D**), and IgG2a (**E**) with ELISA at 1:10 dilution. (**F**) Calculated value of BAL IgG2a/IgG1 ratio. (**G**) Anti-spike total IgG in serum at various dilutions measured by absorbance (A450-630 nm) during ELISA assays. (**H**) Comparison of area under the curve (AUC) of serum anti-spike IgG. (**I**) ACE-2 signal measured by absorbance (A450-630 nm) in spike protein neutralization assay with ELISA. Lower absorbance values indicate higher spike-neutralizing antibody levels in serum. (**J**) Serum from vaccinated mice was assayed for IgG1. (**K**) Comparison of area under the curve (AUC) of serum anti-spike IgG1. (**L**) Serum from vaccinated mice was assayed for IgG2a. (**M**) Comparison of area under the curve (AUC) of serum anti-spike IgG2a. (**N**) Calculated value of serum IgG2a/IgG1 ratio. Error bars represent the SEM. Normality was assessed with the Kolmogorov-Smirnov test. Statistical significance was determined with the Kruskal-Wallis test and Dunn’s post-hoc test for multiple comparisons. *p ≤ 0.05, **p ≤ 0.01, ***p ≤ 0.001, ****p ≤ 0.0001 for all graphs.

Further, we assessed serum systemic IgG and neutralizing antibody (nAb) responses (**Fig. 2, G to I**). The PUUC+CpG PAL-NPs group resulted in significantly increased levels of anti-SARS-CoV-2 IgG in serum at 10^5^-fold dilution (**Fig. 2G**). The serum IgG levels generated (Area under the curve: AUC) after administration of adjuvanated PAL-NPs formulation and control groups follow this order: PUUC+CpG>PUUC>R848+CpG>R848=PAL-NPs (**Fig. 2H**). nAbs play a crucial role in reducing the replication of SARS-CoV-2 and are essential in protecting against severe infections caused by the virus (*45, 46*). We assessed generated serum nAbs by measuring the inhibition of the Spike RBD-ACE2 interaction using an ELISA-based ACE2 competition assay. We found that PUUC+CpG PAL-NPs group elicited high titers of neutralizing antibodies. Spike protein neutralization was detectable at up to a 10^4^-fold dilution in PUUC+CpG PAL-NPs group, which is higher than any other groups (lower absorbance value: A450-A630), and the total neutralization was observed at 100-fold sera dilution (**Fig. 2I**). To evaluate TH1/TH2 skewed IgG response, elicited through IM-Prime/IN-Boost immunization, we assessed the serum IgG subsets (IgG1 and IgG2a) present. We observe that among various PRR agonist formulations on PAL-NP, the PUUC+CpG and PUUC group alone, demonstrated the similar and highest induction of total IgG1 [**Fig. 2, J and K** (IgG1 dilution curve and AUC)]. We also observe that only PUUC+CpG PAL-NP group significantly induces highest anti-spike IgG2a titers [**Fig. 2, L and M** (IgG2a dilution curve and AUC)]. However, the ratio of IgG2a/IgG1 suggests that PUUC+CpG on PAL-NPs show the highest TH1 biased antibody response compared to other adjuvant groups (**Fig. 2N**). Thus, PUUC+CpG PAL-NPs vaccine formulation shows more potent mucosal IgA and IgG levels in BAL fluid and serum. These findings indicate that the multiadjuvant PUUC+CpG PAL-NPs elicit a stronger mucosal antibody (IgA and IgG) response, systemic antibody response, and neutralizing antibody titers, highlighting their significant potential for use in future mucosal subunit nanovaccines.

### PAL-NPs protein subunit vaccine adjuvanated with RIG-I (PUUC) and TLR9 (CpG) agonists induces robust SARS-CoV-2 mucosal T cell and B cell immune responses, when delivered IM-Prime/IN-Boost

The adaptive cellular immune responses were generated and functional in the local tissues during respiratory infection and are responsible for providing long-lasting protective immunity at the infection sites (*16, 47, 48*). Accordingly, we investigated the pulmonary T cell and B cell responses on 35^th^ day (14 days post-boost) of PAL-NPs subunit nanovaccine immunization with IM-Prime/IN-Boost route (**Fig. 3A**). For T cell responses, the single-cell suspension of harvested lungs was restimulated with overlapped spike peptide pools for 6 h and stained with canonical T cell markers and further analyzed with flow cytometry. Memory CD4^+^ and CD8^+^ T cells are more prominent in the local tissues and are non-circulating, known as tissue-resident memory T cells (TRM) (*49*). Traditionally, adjuvants are more responsible for the induction of potent antigen- specific T cell responses in protein subunit vaccines (*31*). The gating strategy for lung T cells subset is shown in supplementary **fig. S8A**. We observed a significantly highest expression of CD4^+^CD69^+^ T cells (∼4.96 fold) in the lung of PUUC+CpG PAL-NP vaccine immunized mice compared to PBS, which is also highest among all other PAL-NP groups, as shown in **Fig. 3, B and C**, with flow cytometry plots (FCM) and the cell percentage. The CD69 is the earliest activation marker generated on the surfaces of antigen-specific activated lymphocytes (*50*). The PUUC PAL-NP vaccine formulation also significantly increases the expression of CD4^+^CD69^+^ T cell population (∼2.95 fold) (**Fig. 3C**). Interestingly, a significant amount of CD4^+^ T cells co- expresses both CD69^+^ and CD103^+^ markers, when mice were vaccinated with PUUC+CpG PAL- NP vaccine formulation (∼2.4 fold), which indicates the presence of lung CD4^+^ TRM responses (**Fig. 3D**).

**Fig. 3.**
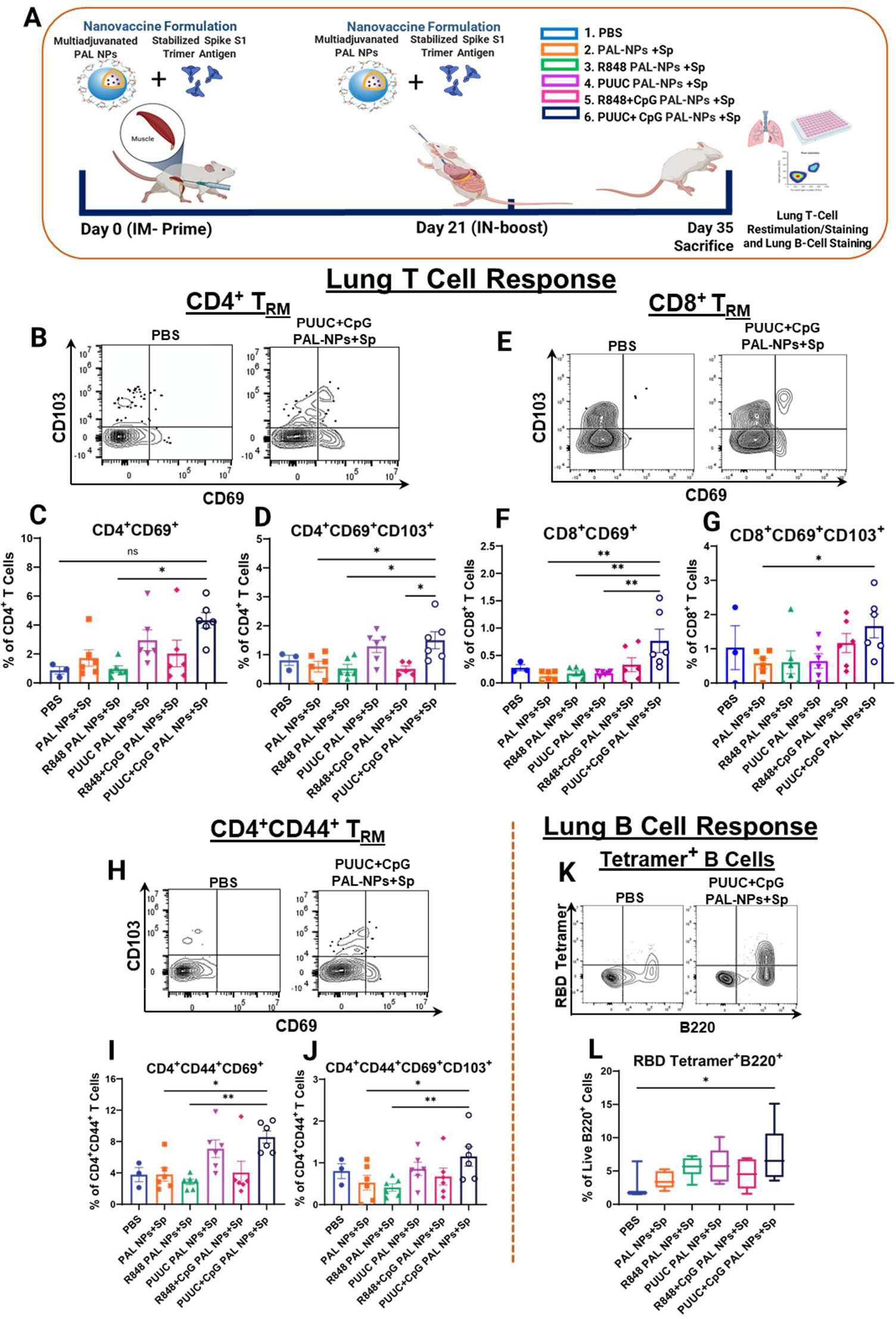
A subunit nanovaccine formulation of PAL-NPs adjuvanated with RIG-I (PUUC) and TLR9 (CpG) agonists, mixed with S1 spike protein elicit robust SARS-CoV-2 mucosal cellular immunity, when delivered IM-Prime/IN-Boost. (**A**) Experimental schematics: Female BALB/c mice (n=3 for PBS and n=6 for adjuvanated PAL-NP formulations) were immunized IM into both anterior tibialis muscles at day 0 (1^st^ dose) with vaccine formulation of adjuvanated PAL- NPs (NPs: 250 µg, PUUC: 20 µg, CpG: 40 µg, and R848: 20 µg) and combined with stabilized spike (Sp) S1 trimer protein at a dose of 1000 ng respectively. On day 21, mice received the 2^nd^dose of protein subunit vaccine formulation IN, using similar doses of adjuvants, PAL-NPs, and protein, except for the CpG dose reduced to 20 µg. Mice were euthanized on day 35 to collect lungs. Lung cells were restimulated with spike peptide pools for 6 h and stained for analysis by flow cytometry. (**B** to **D**) Representative flow cytometry plots (FCM) (**B)** and percentage of CD4^+^CD69^+^ (**C)** and CD4^+^CD69^+^CD103^+^ (**D**) T cell population. (**E** to **G**) Representative FCM plots (**E**) and percentage of CD8^+^CD69^+^ (**F**) and CD8^+^CD69^+^CD103^+^ (**G**) T cell population. (**H** to **J**) Representative FCM plots (**H**) and percentage of CD4^+^CD44^+^CD69^+^ (**I**) and CD4^+^CD44^+^CD69^+^CD103^+^ (**J**) T cell population. Lung cells were stained for B cell markers and analyzed by flow cytometry. (**K** and **L**) Representative FCM plots and percentage of RBD tetramer^+^ B220 cells. Outliers were identified by the ROUT method and removed. Error bars represent the SEM. Statistical significance was calculated using one-way ANOVA followed by Tukey’s post-hoc test for the figures (**C**), (**D**), (**F**), and (**I**), and Bonferroni’s post-hoc test for figure (**G**), (**J**), and (**L**), for multiple comparisons. \**p* ≤ 0.05, \*\**p* ≤ 0.01, \*\*\**p* ≤ 0.001, \*\*\*\**p* ≤ 0.0001 for all graphs. ns represents the non-significant values.

Similar to CD4^+^ cell responses, we further analyzed the CD8^+^ T cell responses after PAL- NP vaccine immunization. We observed that lung CD8^+^ T cells significantly expressed substantially higher levels of CD69^+^ activation marker (∼2.15-fold), [**Fig. 3, E and F** (FCM plots and cell percentage)] and CD69^+^CD103^+^ tissue-resident memory markers (∼1.4 fold), [**Fig. 3, E and G** (FCM plots and cell percentage)], after PUUC+CpG PAL-NP vaccine immunization. We did not observe the significant population of effector memory CD4^+^CD44^+^ T cells (**fig. S3, A and E**), but further gating shows the significant expression of CD4^+^CD44^+^CD69^+^ and CD4^+^CD44^+^CD69^+^CD103^+^ (resident memory) cell population with both PUUC+CpG and PUUC PAL-NP vaccine group and comparatively higher in PUUC+CpG PAL-NP group (**Fig. 3, H to J**). This population represents the effector resident memory T cell population (Eff TRM). We also evaluated the TRM cell population from total T cells (CD3^+^), which is significantly higher with the PUUC+CpG PAL-NP vaccine formulation and correlates with high frequencies of individual CD4^+^ and CD8^+^ TRM cell population (**fig. S2, A and B**). With the same formulation, a double negative CD4^-^CD8^-^ population was also observed in T cell subset, which could be γδ T cells, probably activated in an MHC-independent manner and found in epithelial and mucosal tissues (**fig. S3K**). This result shows that PUUC+CpG PAL-NP vaccine formulation induces strong local antiviral T cell immunity when administered IM-prime and IN-boost in mice.

The presence of anti-viral memory B cell responses in the local tissues will also contribute to long-term immune protection against SARS-CoV-2 (*51*). Therefore, we examined induced SARS-CoV-2 specific B cell responses with adjuvanated PAL-NP vaccine formulations via IM-Prime/IN-Boost immunization. After lung harvesting, single-cell suspension was stained with different B cell markers: antigen (RBD) specific, antibody-secreting cells (ASC), GC B cells, and memory B cells (tissue-resident memory B cells: BRM), and analyzed with the flow cytometry. The gating strategy for lung B cells is shown in supplementary **fig. S8B**. Mice immunized with PUUC+CpG PAL-NP vaccine formulation through IM-Prime/IN-Boost route show a significant RBD tetramer^+^ B cell population [**Fig. 3, K and L** (FCM plots and percentage)], which is specific for receptor binding domain (RBD) of the spike protein. Compared to PBS control, the RBD tetramer+ B cells are two-fold higher in mice immunized with PUUC+CpG PAL-NP vaccine formulation. In addition, upon IM-Prime/IN-Boost administration of PUUC+CpG PAL-NP vaccine formulation, we have observed the presence (non-significant) of several types of immune cells, including antibody-secreting cells (ASC: B220^+/-^CD138^+^), IgG^+^ antibody-secreting cells (IgG^+^ASC: B220^+/-^CD138^+^IgA^-^), IgM^+^ memory B cells (B220^+^IgM^+^IgD^-^CD38^+^), and IgG^+^ BRM (B220^+^IgD^-^IgM^-^CD38^+^IgA^-^), as compared to control groups and other adjuvant groups (**figs. S4A, S4C, S4E, S4G, and S4H**). Interestingly, R848+CpG PAL-NP vaccine immunized mice with IM-Prime/IN-Boost strategy show a non-significant IgA^+^ ASC (IgA^+^ASC: B220^+/-^CD138^+^IgA^+^) and IgA^+^ BRM (B220^+^IgD^-^IgM^-^CD38^+^IgA^+^) cell population (**fig. S4B and S4D**).

### PUUC+CpG and PUUC PAL-NP subunit vaccine formulation enhances TH1 type immunity and reduces the secretion of TH2 type cytokines, when delivered IM-Prime/IN-Boost

To further investigate the TH1/TH2 expression profile of the T cell population, restimulated lung cells from spike peptide pool were stained with intracellular cytokines (Tumor necrosis factor-alpha: TNF-α, Interferon-gamma: IFN-γ, and Granzyme B: GrzB) and analyzed with flow cytometry (**Fig. 4A**). Mice immunized with PUUC+CpG PAL-NP vaccine formulation significantly increased the frequency of monofunctional CD4^+^ TRM which are enriched for TH1 type cytokine TNF-α (**Fig. 4, B and C**). PUUC PAL-NP vaccine formulation also shows a significant increase in the frequency of monofunctional CD4^+^ TRM population in lungs that express TH1 type cytokine IFN-γ (**Fig. 4, D and E**). Similar to CD4^+^ TRM, the induction of CD8^+^ TRM cell population expressed TNF-α, is also enhanced by PUUC+CpG PAL-NP immunization (**Fig. 4, F and G**). Lungs from mice immunized with PUUC+CpG PAL-NP vaccine formulation had a non-significant IFN-γ enriched CD8^+^ TRM cell population (**Fig. 4, H and I**). The PUUC PAL-NP group enhances the frequency of CD4^+^CD44^+^ TRM cell population that expresses GrzB (*p*=0.0564) (**Fig. 4, J and K**). The PUUC PAL-NP vaccine formulation alone, also increases the percentage of GrzB expressing monofunctional CD4^+^CD44^+^ TRM, and CD4^+^CD44^+^ T cell population in a non-significant manner [**Fig. 4, L and M, and fig. S3D**]. PUUC+CpG PAL-NP vaccine formulation significantly increases the percentage of GrzB expressing monofunctional CD8^+^CD44^+^ T cell population in lungs (**fig. S3H**). A significant population of CD4^+^ TRM cells enriched with GrzB was observed in the mice vaccinated with PUUC PAL-NPs formulation (**fig. S2I**). Whereas both PUUC+CpG and PUUC PAL-NP formulations did not increase a significant population of GrzB expressing CD8^+^ TRM cells in the mice lungs (**fig. S2M**). Mice vaccinated with PUUC PAL-NP vaccine formulation have non-significant CD4^+^ TRM polyfunctional cells that co-express TNF-α, and IFN-γ (**fig. S3I**). Similarly, a non-significant increase in the percentage of CD8^+^ TRM polyfunctional cells that co-express both TNF-α and IFN-γ was observed in the lungs of mice vaccinated with PUUC+CpG PAL-NP formulation (**fig. S3J**). PUUC+CpG and R848+CpG PAL- NP vaccine formulations significantly increase the CD4^+^, CD8^+^, CD4^+^ CD44^+^, and CD8^+^CD44^+^cell populations in the mice lungs that express TNF-α (**figs. S2F, S2J, S3B, and S3F**). PUUC+CpG PAL-NP vaccine formulation significantly increased the CD3^+^ TRM that expresses TNF-α (**fig. S2C**), whereas the PUUC PAL-NP group showed a non-significant increase in total CD3^+^ TRM that expresses IFN-γ and GrzB (**fig. S2, D and E**). PUUC+CpG PAL-NP group induces a significant increase in the GrzB expressing monofunctional CD8^+^ TRM cell population, but PUUC PAL-NP group induces a non-significant increase in GrzB expressing monofunctional CD4^+^ TRM cell population (**fig. S2, G and K**). None of the adjuvanted PAL-NP formulations increase the CD4^+^ IFN-γ T cell population in the mice lungs (**fig. S2, H and L**). For a more comprehensive study of TH1/TH2 cytokine profile, we performed a multiplexed cytokine assay to assess various cytokine concentrations from supernatants of lung T cells after restimulation. Secreted cytokine profile is associated more with the TH1 type response, where PUUC+CpG PAL-NP vaccinated mice secrete TH1 type cytokine TNF-α (**Fig. 4O**), and PUUC PAL-NP vaccinated mice secrete TH1 type cytokine IFN-γ (**Fig. 4N**). This profile is consistent with prior assessment by flow cytometry data, where TNF-α and IFN-γ expressing T cells were significantly higher in RIG-I adjuvanated PAL-NPs (PUUC+CpG) than the cohorts immunized with other adjuvants and controls. Thus, the elevated level of TH1 type (TNF-α and IFN-γ) cytokine (**Fig. 4, N and O**) and suppressed (IL-10, IL-4) (**Fig. 4, P and Q**) or very low detection level of TH2 type cytokine (IL- 13) profile were observed in PUUC+CpG PAL-NP vaccinated mice (**fig. S3L**). These results demonstrate that RIG-I targeted PAL-NP combination greatly enhanced T cells’ potency and TH1 biased response.

**Fig. 4.**
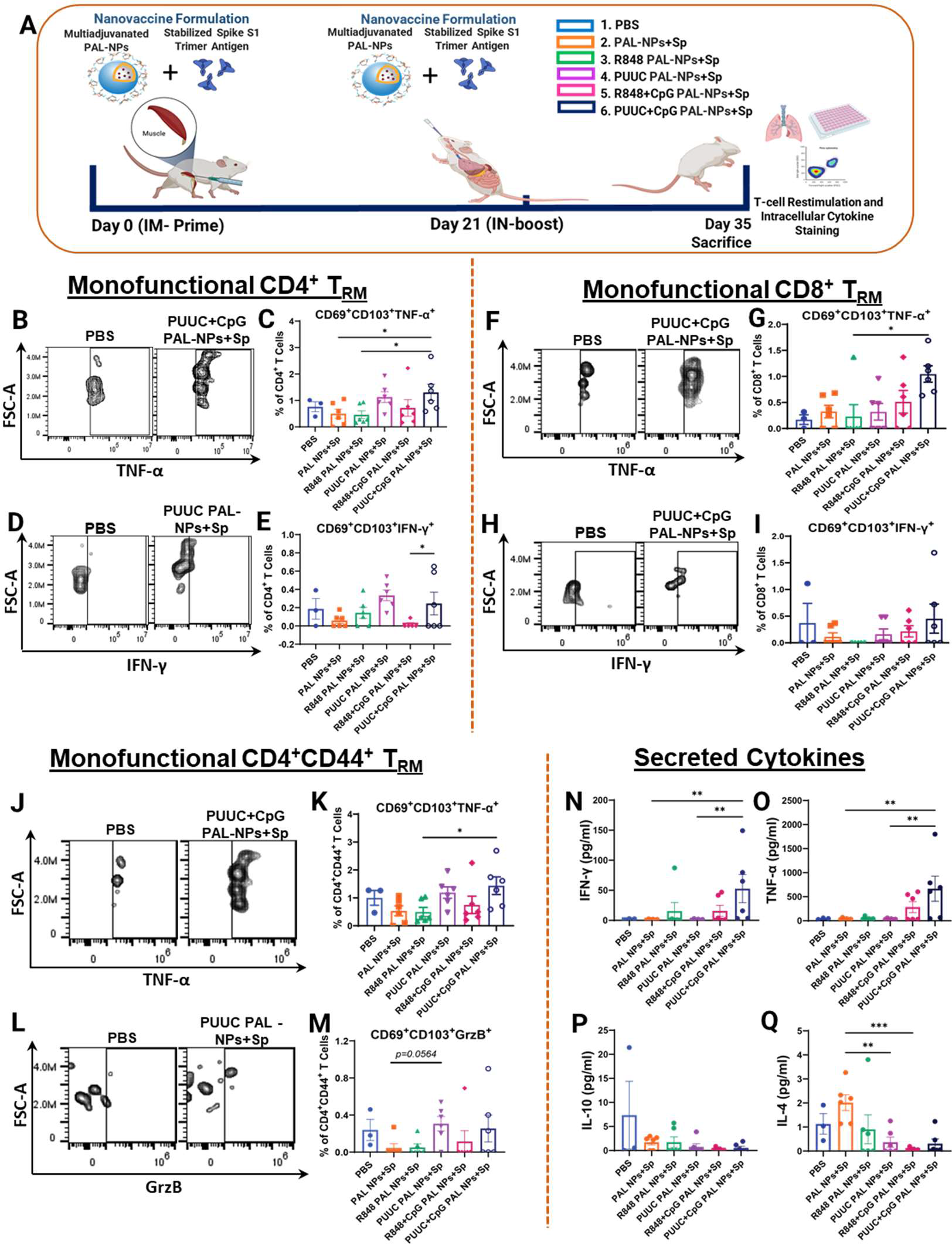
A subunit nanovaccine formulation of PAL-NPs adjuvanated with RIG-I (PUUC) and TLR9 (CpG) agonists, mixed with S1 spike protein elicit robust SARS-CoV-2 mucosal cellular immunity, when delivered IM-Prime/IN-Boost. (**A**) Experimental schematics: Female BALB/c mice (n=3 for PBS and n=6 for other formulations) were immunized IM into both anterior tibialis muscles at day 0 (1^st^ dose) with vaccine formulation of multiadjuvanated PAL-NPs (NPs: 250 µg, PUUC: 20 µg, CpG: 40 µg, and R848: 20 µg) and combined with stabilized spike (Sp) S1 trimer protein at a dose of 1000 ng respectively. On day 21, mice received the 2^nd^ dose of protein subunit vaccine formulation IN, using similar doses of adjuvants, PAL-NPs, and protein, except for the CpG dose reduced to 20 µg. Mice were euthanized on day 35 to collect lungs. Lung cells were restimulated with spike peptide pools for 6 h and stained for analysis by flow cytometry. (**B** and **C**) Representative flow cytometry plots (FCM) of monofunctional CD4^+^ TRM and percentages of cells expressing TNF-α. (**D** and **E**) Representative FCM plots of monofunctional CD4^+^ TRM and percentages of cells expressing IFN-γ. (**F** and **G**) Representative FCM plots of monofunctional CD8^+^ TRM and percentages of cells expressing TNF-α. (**H** and **I**) Representative FCM plots of monofunctional CD8^+^ TRM and percentages of cells expressing IFN-γ. (**J** and **K**) Representative FCM plots of monofunctional CD4^+^CD44^+^ TRM and percentages of cells expressing TNF-α. (**L** and **M**) Representative FCM plots of monofunctional CD4^+^CD44^+^ TRM and percentages of cells expressing GrzB. **(N** to **Q)** Cytokine concentration in supernatants from restimulated lung cells TNF-α (**N**), IFN-γ (**O**), IL-10 (**P**), IL-4 (**Q**). Error bars represent the SEM. Outliers were identified by the ROUT method and removed. Statistical significance was calculated using one-way ANOVA followed by Tukey’s post-hoc test for figures (**I**) and (**K** to **Q**), and Bonferroni’s post-hoc test for figures (**C**), (**E**), and (**G**), for multiple comparisons. \**p* ≤ 0.05, \*\**p* ≤ 0.01, \*\*\**p* ≤ 0.001, \*\*\*\**p* ≤ 0.0001 for all graphs. ns represents the non-significant values.

### PUUC+CpG PAL-NPs subunit vaccine formulation elicits comparable SARS-CoV-2 mucosal humoral immunity with IN-Prime/IN-Boost

PUUC+CpG PAL-NP vaccine formulation was found to be one of the highly promising candidates that shows strong mucosal humoral and local cellular and, systemic responses with IM-Prime/IN-Boost. As a result, we conducted a more comprehensive second in vivo study, which is more concentrated on immune responses specific to the prime-boost vaccine administration routes. PUUC+CpG PAL-NP vaccine formulation is administered in mice with three prime-boost strategies: (A) IM-Prime/IN-Boost, (B) IN-Prime/IM-Boost, and (C) IN-Prime/IN-Boost. Mice were immunized prime at day 0 and boosted at day 21 with PUUC+CpG PAL-NP vaccine formulation and sacrificed at day 35. BAL fluid and serum samples were collected to analyze generated mucosal and systemic humoral response (**Fig. 5A**). We first evaluated mucosal anti-spike S1 IgA, IgG, and neutralizing antibody responses generated in BAL fluid through these three prime-boost strategies (**Fig. 5, B to F**). Consistent with the first in vivo study, the IM-Prime/IN-Boost route enhances mucosal and systemic antibody responses. Surprisingly, IN-Boosting with PUUC+CpG PAL-NP vaccine formulation after IN-Priming (IN-Prime/IN-Boost) induces substantial and significant anti-spike IgA level (A450-A630 = 2.4) in BAL fluid (at 1:2 dilution) (**Fig. 5B**). Interestingly, a similar level of strong nAbs were induced with both IM-Prime/IN-Boost and IN-Prime/IN-Boost routes (at 1:2 dilution) (**Fig. 5D**). However, the total IgG level in BAL fluid is significantly higher in IM-Prime/IN-Boost, and follow this order: IM-Prime/IN-Boost> IN-Prime/IM-Boost> IN-Prime/IN-Boost (**Fig. 5C**). The IM-Prime/IN-Boost and IN-Prime/IM- Boost groups show more TH1 type immunity (IgG2a/IgG1 ratio>1), whereas the IN-Prime/IN-Boost group is close to IgG2a biased immunity (**Fig. 5, E and F, and fig. S5C**). The systemic humoral immune responses generated through IM-Prime/IN-Boost route are consistent with the first *in vivo* study results, showing a high and significant level of total IgG and nAb levels (**Fig. 5, G to I**). The secreted total IgG (AUC value) levels in serum after the PUUC+CpG PAL-NP immunization in mice with three prime-boost strategies follow the order: IM-Prime/IN-Boost>IN- Prime/IM-Boost>IN-Prime/IN-Boost [**Fig. 5, G and H** (IgG dilution curve and AUC)]. Interestingly, both IM-Prime/IN-Boost and IN-Prime/IM-Boost groups show more TH1 type immunity, whereas IN-Prime/IN-Boost group is close to IgG2a biased immunity [**Fig. 5, J and K** (IgG1 dilution curve and AUC), and **Fig. 5, L and M** (IgG2a dilution curve and AUC)]. With IN- Prime/IN-Boost, a significant serum IgA level group was observed, but it is comparatively lower than IM-Prime/IN-Boost group and follows the order IM-Prime/IN-Boost >IN-Prime/IN-Boost> IN-Prime/IM-Boost [(**Fig. 5, N and O** (IgA dilution curve and AUC)]. These results altogether indicate that the PUUC+CpG PAL-NP vaccine formulation generates more potent mucosal and systemic humoral responses with the IM-Prime/IN-Boost group. However, the induction of strong mucosal IgA (comparable to IM-Prime/IN-Boost) and nAb level (similar to IM-Prime/IN-Boost) in the BAL fluid with the IN-Prime/IN-Boost group make this PUUC+CpG PAL-NP vaccine formulation also suitable for fully mucosal SARS-CoV-2 vaccine.

**Fig. 5.**
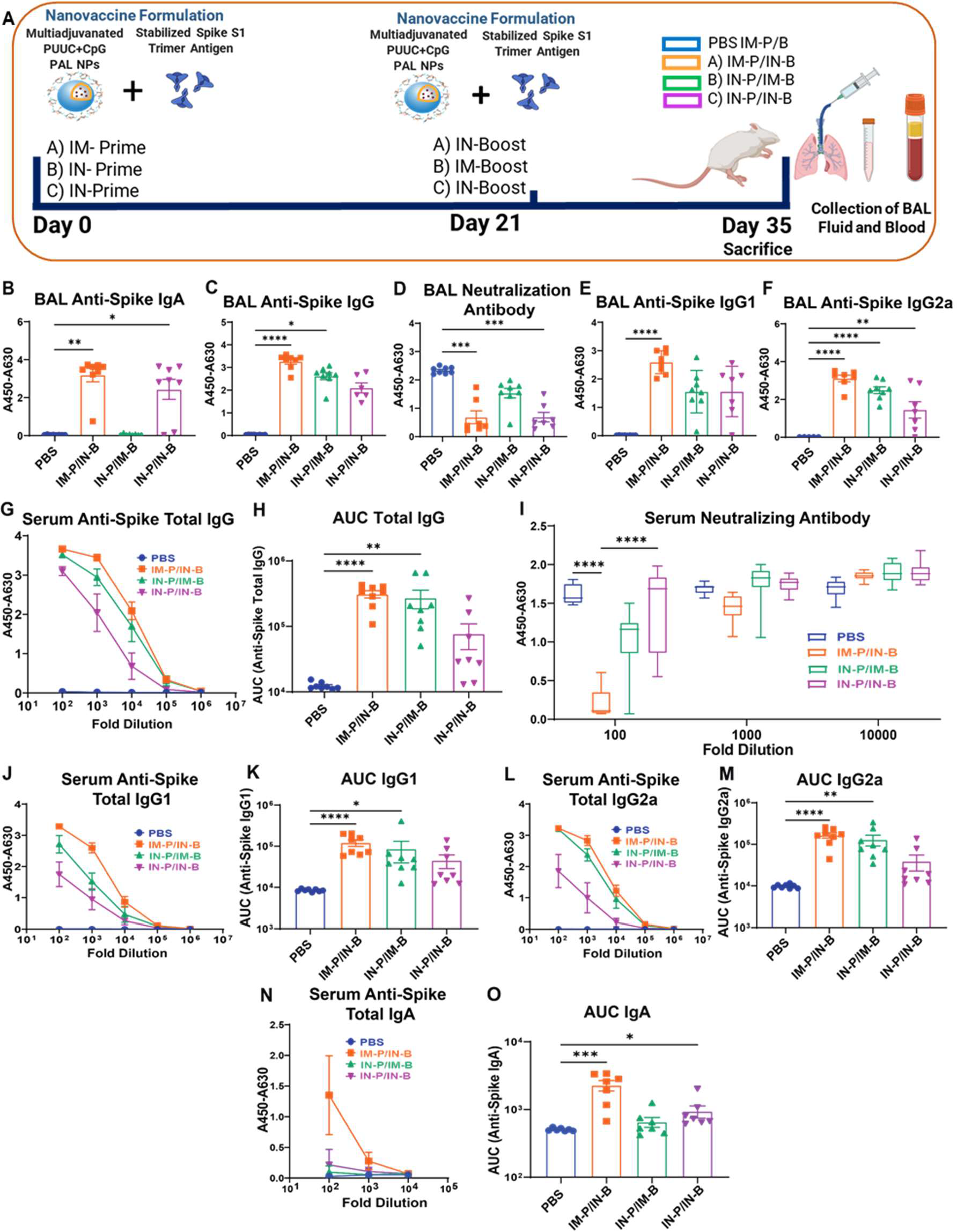
PUUC+CpG PAL-NPs protein subunit vaccine formulation, elicit robust SARS-CoV- 2 mucosal and systemic humoral immunity with IM-Prime/IN-Boost group and induces a significant level of mucosal humoral responses with IN-Prime/IN-Boost group. (**A**) Experimental Schematics: Female BALB/c mice (n=8 for all groups) were immunized with three prime-boost strategies. At day 0 (1^st^ dose), vaccine formulation of PUUC+CpG PAL-NPs (NPs: 250 µg, PUUC: 20 µg, CpG: 40 µg, and R848: 20 µg) combined with stabilized spike (Sp) S1 trimer protein (1000 ng) was administered. On day 21, mice received the 2^nd^ dose of protein subunit vaccine formulation IN (CpG dose reduced to 20 µg). Mice were euthanized on day 35 to collect BAL fluid, and blood. BAL and serum from vaccinated mice was assayed with ELISA. (**B** to **F**) BAL fluid from vaccinated mice was assayed for anti-spike IgA (**B**), IgG (**C**), spike neutralization antibody (**D**), IgG1 (**E**), and IgG2a (**F**) with ELISA at 1:5 dilution except for IgA and neutralization assay which was performed at 1:2 dilution. (**G**) Anti-spike total IgG in serum at various dilutions measured by absorbance (A450-630 nm) during ELISA assay. (**H**) Comparison of area under the curve (AUC) of serum anti-spike IgG. (**I**) ACE-2 signal measured by absorbance at 450 nm in spike protein neutralization assay with ELISA. (**J**) Serum from vaccinated mice was assayed for IgG1. (**K**) Comparison of area under the curve (AUC) of serum anti-spike IgG1. (**L**) Serum from vaccinated mice was assayed for IgG2a. (**M**) Comparison of area under the curve (AUC) of serum anti-spike IgG2a. (**N**) Serum from vaccinated mice was assayed for IgA. (**O**) Comparison of area under the curve (AUC) of serum anti-spike IgA. Error bars represent the SEM. Normality was assessed with the Kolmogorov-Smirnov test. Statistical significance was determined with the Kruskal-Wallis test and Dunn’s post-hoc test for multiple comparisons. *p ≤ 0.05, **p ≤ 0.01, ***p ≤ 0.001, ****p ≤ 0.0001 for all graphs. ns represent the not significant values.

### PUUC+CpG PAL-NP subunit vaccine elicits robust SARS-CoV-2 T cell (Tissue-resident memory) immunity with IN-Prime/IN-Boost and B cell responses with IM-Prime/IN-Boost

In the second in vivo study, we further investigate route-specific cellular immune responses (T cell and B cell) with three prime-boost strategies: (A) IM-Prime/IN-Boost, (B) IN-Prime/IM-Boost, and (C) IN-Prime/IN-Boost) using PUUC+CpG PAL-NPs subunit vaccine formulation. For T cell responses, the lung single-cell suspension of harvested lungs was restimulated with spike peptide pools for 6 h, stained with canonical T cell markers, and further analyzed with flow cytometry (**Fig. 6A**). Gating strategy for lung T cells is shown in supplementary **fig. S8A**. PUUC+CpG PAL-NPs group induced stronger and enhanced local T cell responses when delivered IN-Prime/IN-Boost, which are surprisingly higher than IM-Prime/IN-Boost. Analyzing tissue-resident memory T cells (TRM), we observed that a proportion of CD4^+^ helper T cells significantly express both CD69^+^ and CD69^+^CD103^+^ (CD4^+^ TRM) cells with IM-Prime/IN-Boost and IN-Prime/IN-Boost group, but the population frequency is higher in the later group [**Fig. 6, B and C** (FCM plots and CD4^+^CD69^+^ T cell percentage), and **Fig. 6D** (CD4^+^ TRM percentage)]. The frequency of CD4^+^CD69^+^ cells is also higher in IN-Prime/IN-Boost group (∼2.73 fold) than IM-Prime/IN-Boost group (∼2.28 fold) with respect to the PBS control. Similarly, the frequency of CD4^+^CD69^+^CD103^+^ population is slightly higher in IN-Prime/IN-Boost group (∼2.95 fold) than IM-Prime/IN-Boost group (∼2.92 fold). Overall, the frequency of generated CD4^+^CD69^+^ T cell population follows the order: IN-Prime/IN-Boost>IM-Prime/IN-Boost>IN-Prime/IM-Boost. In comparison, the CD4^+^CD69^+^CD103^+^ T cell population frequency follows the order: IN-Prime/IN- Prime>IM-Prime/IN-Boost>IN-Prime/IM-Boost.

**Fig. 6.**
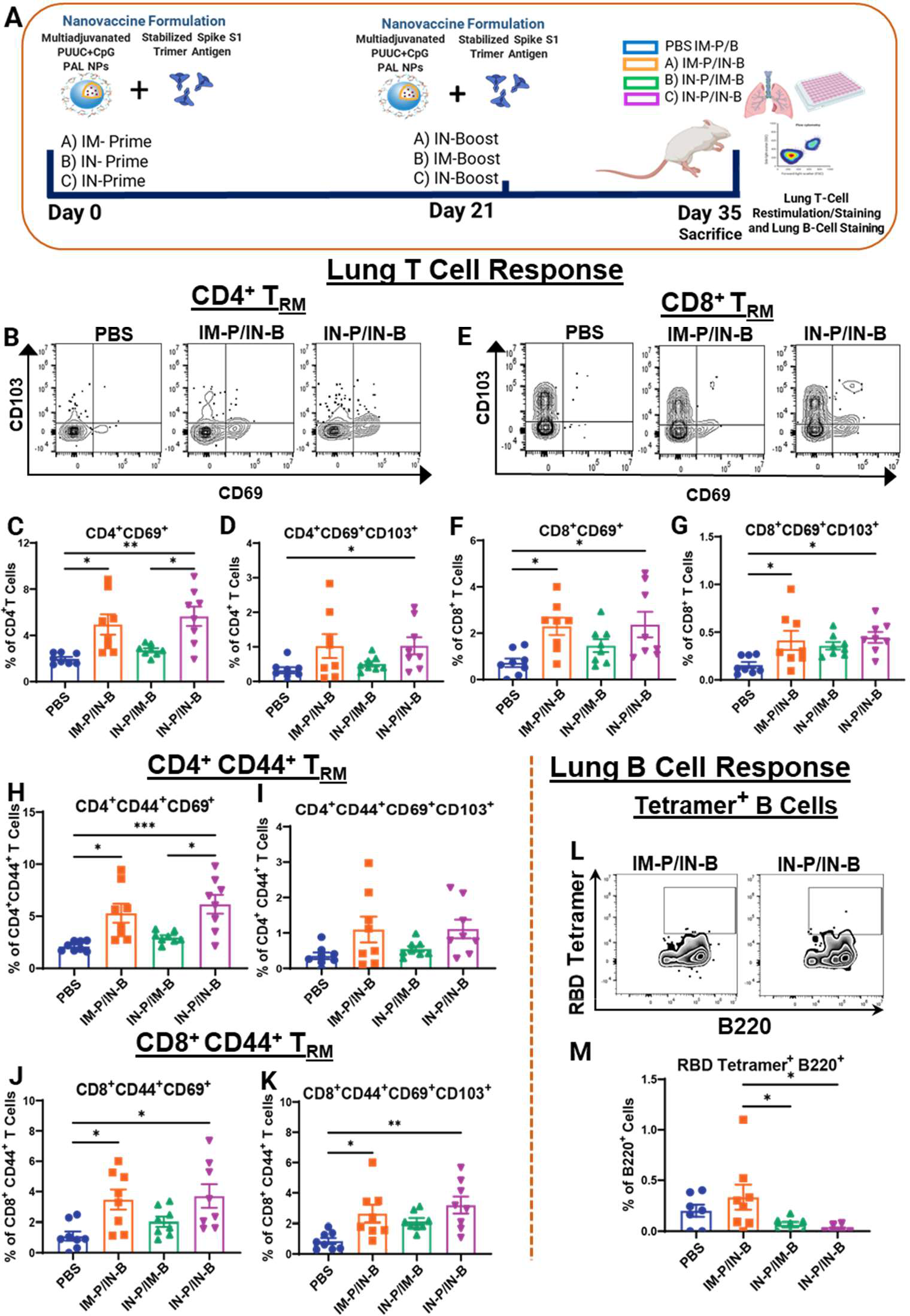
PUUC+CpG PAL-NPs protein subunit vaccine formulation elicits robust SARS-CoV- 2 T cell (TRM) immunity with IN-Prime/IN-Boost and B cell responses with IM-Prime/IN- Boost. (**A**) Experimental schematics: Female BALB/c mice (n=8 for all groups) were immunized with three prime-boost strategies. At day 0 (1^st^ dose), a vaccine formulation of PUUC+CpG PAL-NPs combined with stabilized spike protein (Sp) S1 trimer protein was administered. On day 21, mice received the 2^nd^ dose of protein subunit vaccine formulation IN (CpG dose reduced to 20 µg). Mice were euthanized after two weeks on day 35 to collect lungs. Lung cells were restimulated with spike peptide pools for 6 h and stained for analysis by flow cytometry. (**B** to **D**) Representative flow cytometry plots (FCM groups: PBS, IM-Prime/IN-Boost, and IN-Prime/IN-Boost) (**B**) and percentage of CD4^+^CD69^+^ (**C**) and CD4^+^ TRM (**D**) cell population. (**E** to **G**) Representative FCM plots (groups: PBS, IM-Prime/IN-Boost, and IN-Prime/IN-Boost) (**E**) and percentage of CD8^+^CD69^+^ (**F**) and CD8^+^ TRM (**G**) cell population. Percentage of T cell populations: CD4^+^CD44^+^CD69^+^ (**H**), CD4^+^CD44^+^ TRM (**I**), CD4^+^CD44^+^CD69^+^ (**J**), CD4^+^CD44^+^ TRM (**K**). (**L** and **M**) Representative flow cytometry plots and percentage of RBD tetramer^+^ B220^+^ cells. Error bars represent the SEM. Statistical significance was calculated with One-Way ANOVA and Tukey post-hoc test for multiple comparisons. \**p* ≤ 0.05, \*\**p* ≤ 0.01, \*\*\**p* ≤ 0.001, \*\*\*\**p* ≤ 0.0001 for all graphs. ns represents the not significant values.

With CD8^+^ T cells responses, we observed similar results. The frequency of CD8^+^CD69^+^ T cells is higher in the mice immunized through IN-Prime/IN-Boost strategy (∼2.37 fold) compared to IM-Prime/IN-Boost (∼2.29 fold) [**Fig. 6, E and F** (FCM plots and percentage)]. The frequency of CD8^+^ TRM in the IN-Prime/IN-Boost group (∼0.43 fold) is close to IM-Prime/IN-Boost group (∼0.4 fold) [**Fig. 6, E and G** (FCM plots and percentage)]. We also found that a proportion of CD4^+^CD44^+^ cells shows both CD69^+^ and CD69^+^CD103^+^ populations, with both IN- Prime/IN-Boost and IM-Prime/IN-Boost groups, which indicate the presence of effector memory resident cell population (Eff TRM) (**Fig. 6, H and I**). However, the frequency of CD4^+^CD44^+^CD69^+^ cells are significantly higher in the IN-Prime/IN-Boost group, and the frequency of CD4^+^CD44^+^CD69^+^CD103^+^ cells is almost similar in both IN-Prime/IN-Boost and IM-Prime/IN-Boost groups. Like the CD4^+^CD44^+^ T cell response, we analyzed the CD8^+^CD44^+^ T cell responses after immunization, and similar results were observed. The frequencies of CD8^+^CD44^+^CD69^+^ and CD8^+^CD44^+^CD69^+^CD103^+^ T cells were also comparatively higher in IN-Prime/IN-Boost than IM-Prime/IN-Boost group. (**Fig. 6, J and K**). We also evaluated the TRM cell population from total T cells (CD3^+^), which is higher with the IN-Prime/IN-Boost route and correlates with the frequencies of individual CD4^+^ and CD8^+^ TRM cell populations (**fig. S5A and S5B**). Further confirmation of γδ phenotype from the first in vivo study, the lung cells were stained with TCRγδ^+^ marker in the second in vivo study. We observed a high CD3^+^CD4^-^CD8^-^TCRγδ^+^ cell population with IN-Prime/IN-Boost group (**fig. S5N**). These results show that IN-Prime/IN-Boost administration of PUUC+CpG PAL-NPs vaccine formulation induces strong antiviral T cell immunity in the lungs.

In addition to antiviral memory T cell responses, B cell responses are also necessary for clearing mucosal pathogens (*51*) Therefore, in the second in vivo study, we examined B cell responses generated with three prime-boost strategies (A) IM-Prime/IN-Boost, (B) IN-Prime/IM- Boost, and (C) IN-Prime/IN-Boost) using PUUC+CpG PAL-NPs vaccine formulation (gating strategy: supplementary **fig. S8B**). We observed a significantly higher RBD tetramer^+^ B cell population with IM-Prime/IN-Boost, compared to IN-Prime/IM-Boost and IN-Prime/IN-Boost groups, which is specific for receptor binding domain (RBD) of spike protein [**Fig. 6, L and M** (FCM plots and percentage)]. At the same time, a non-significant increment in IgA^+^ antibody-secreting cells (IgA^+^ASC: B220^+/-^CD138^+^IgA^+^) was observed with IM-Prime/IN-Boost route (**fig. S7B**). The presence of IgA^+^ tissue-resident memory B cells (IgA^+^BRM: B220^+^IgD^-^IgM^-^CD38^+^IgA^+^), GC-B cells (B220^+^CD38^-^GL7^+^), and IgM^+^ Memory B cells (B220^+^IgD^-^IgM^+^CD38^+^) was also observed in a non-significant manner with IN-Prime/IN-Boost group (**figs. S7E, S7D, and S7C**).

### PUUC+CpG PAL-NPs subunit vaccine formulation enhances TH1 type immunity with IN- Prime/IN-Boost group and reduces the secretion of TH2 type cytokines

In the second in vivo study, we further examine TH1/TH2 expression profile by administrating the PUUC+CpG PAL-NP vaccine formulation in mice with three prime-boost strategies: (A) IM-Prime/IN-Boost, (B) IN-Prime/IM-Boost, and (C) IN-Prime/IN-Boost (**Fig. 7A**). After harvesting of lungs, the restimulated T cells from spike peptide pool, was further stained with intracellular cytokines: TNF-α, IFN-γ, and GrzB, and analyzed with flow cytometry. With both IN-Prime/IN-Boost and IM-Prime/IN-Boost groups, we observed a significant monofunctional CD4^+^ TRM population that expresses TH1 type intracellular cytokines: TNF-α, IFN-γ and cytotoxic GrzB [**Fig. 7, B to D** (FCM plots), and **Fig. 7E** (percentage)]. Similarly, with both the IN-Prime/IN-Boost and IM-Prime/IN-Boost groups, we observed a significant monofunctional CD8^+^ TRM population that expresses TH1 type intracellular cytokines: TNF-α, IFN- γ, and cytotoxic GrzB [(**Fig. 7, F to H** (FCM plots), and **Fig. 7I** (percentage)]. We also observed the higher polyfunctional CD4^+^ TRM cell population, which co-express TH1 type cytokines: TNF- α and GrzB (**Fig. 7J**), IFN-γ and GrzB (**Fig. 7K**), and TNF-α and IFN-γ (**fig. S5O**), with IN- Prime/IN-Boost group compare to IM-Prime/IN-Boost group. Similar results were observed with the CD8^+^ TRM cell population with PUUC+CpG PAL-NP vaccine formulation when immunized IN-Prime/In-Boost (**Fig. 7, L and M and, fig. S5P**). Furthermore, the monofunctional and polyfunctional CD4^+^ TRM and CD8^+^ TRM cells are significantly higher in IN-Prime/IN-Boost group than IM-Prime/IN-Boost group. The IN-Prime/IN-Boost group also induces CD3^+^ TRM cell populations that express TNF-α, IFN-γ, and GrzB (**figs. S5E, S5F, and S5G**). We also observed significantly high monofunctional CD4^+^, CD4^+^CD44^+^, CD8^+^CD44^+^, and CD8^+^ T cell populations enriched for GrzB, with IN-Prime/IN-Boost group (**figs. S5H, S5K, S6B, and fig. S6F**).

**Fig. 7.**
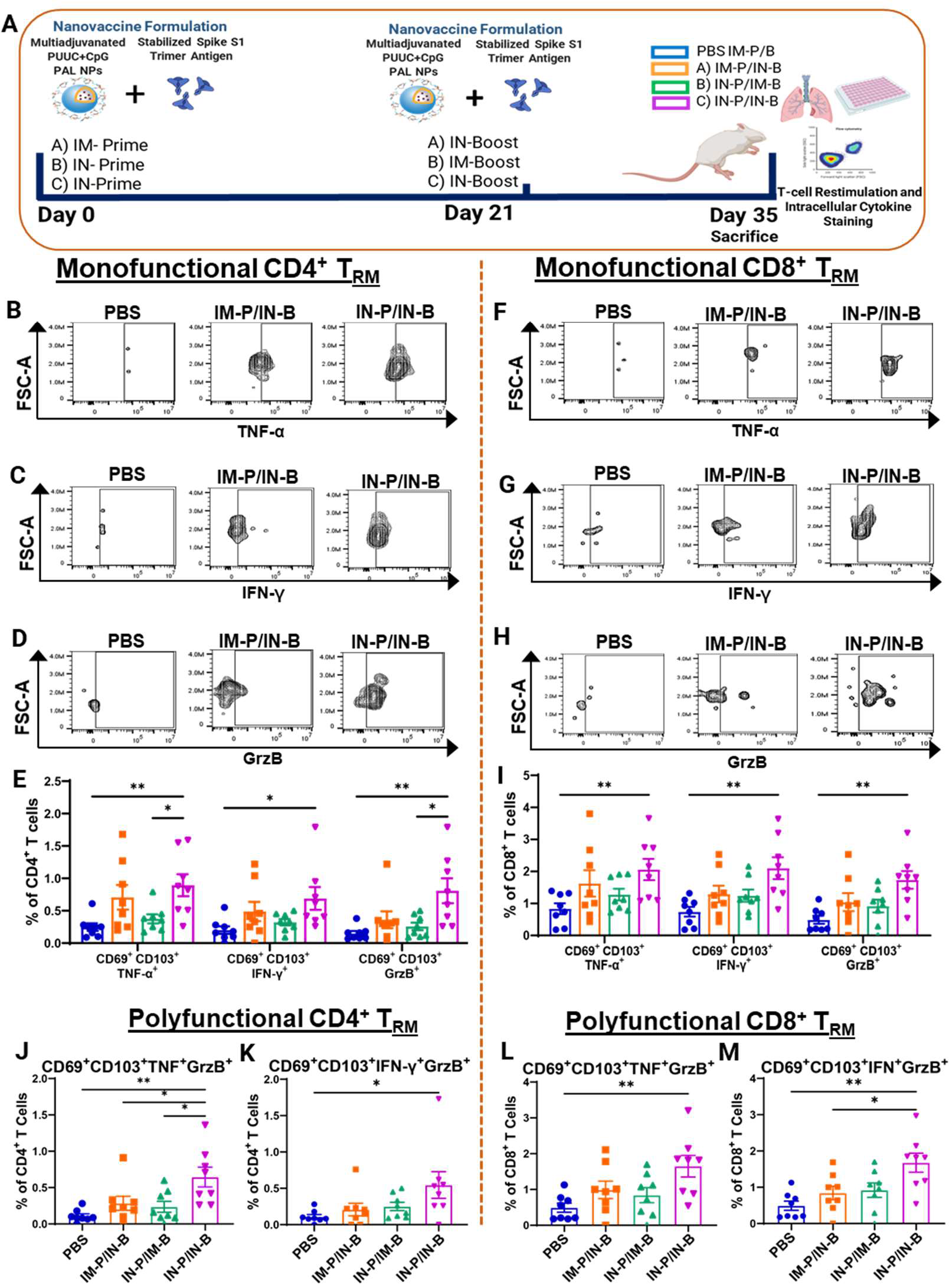
**PUUC+CpG PAL-NPs subunit vaccine formulation enhances TH1 type immunity with IN-Prime/IN-Boost group**. (**A**) Experimental schematics: Female BALB/c mice (n=8 for all groups) were immunized with three prime-boost strategies. At day 0 (1^st^ dose), a vaccine formulation of PUUC+CpG PAL-NPs combined with a stabilized spike (Sp) S1 trimer protein was administered. On day 21, mice received the 2^nd^ dose of protein subunit vaccine formulation IN (CpG dose reduced to 20 µg). Mice were euthanized on day 35 to collect lungs. Lung cells were restimulated with spike peptide pools for 6 h and stained for analysis by flow cytometry. (**B** to **D**) Representative FCM plots (groups: PBS, IM-Prime/IN-Boost, and IN-Prime/IN-Boost) of monofunctional CD4^+^ TRM cells expressing TNF-α (**B**), IFN-γ (**C**), and GrzB (**D**). (**E**) Percentages of monofunctional CD4^+^ TRM cells expressing TNF-α, IFN-γ, and GrzB. (**F** to **H**) Representative FCM plots (groups: PBS, IM-Prime/IN-Boost, and IN-Prime/IN-Boost) of monofunctional CD8^+^ TRM cells expressing TNF-α (**F**), IFN-γ (**G**), and GrzB (**H**). (**I**) Percentages of monofunctional CD8^+^ TRM cells expressing TNF-α, IFN-γ, and GrzB. (**J**) Percentages of polyfunctional CD4^+^ TRM cells co-expressing TNF-α and GrzB. (**K**) Percentages of polyfunctional CD4^+^ TRM cells co- expressing IFN-γ and GrzB. (**L**) Percentages of polyfunctional CD8^+^ TRM cells co-expressing TNF- α and GrzB. (**M**) Percentages of polyfunctional CD8^+^ TRM cells co-expressing IFN-γ and GrzB. Error bars represent the SEM. Statistical significance for cytokine^+^ T cell frequencies was calculated with One-Way ANOVA and Tukey post-hoc test for multiple comparisons. \**p* ≤ 0.05, \*\**p* ≤ 0.01, \*\*\**p* ≤ 0.001, \*\*\*\**p* ≤ 0.0001 for all graphs. ns represent the not significant values.

The IN-Prime/IM-Boost and IN-Prime/IN-Boost groups non-significantly increase the monofunctional CD4^+^, CD4^+^CD44^+^, CD8^+^CD44^+^, and CD8^+^ T cell populations enriched for TNF- α (**figs. S5J, S5M, S6D, and fig. S6H**). There is a non-significant increase in the IFN-γ enriched CD8^+^CD44^+^ cell population with IN-Prime/IN-Boost and IN-Prime/IM-Boost groups (**fig. S6G**). At the same time, the IN-Prime/IN-Boost group shows a significant increase in the frequency of monofunctional CD4^+^CD44^+^ TRM and CD8^+^CD44^+^ TRM cell population that expresses TH1 type cytokines: TNF-α, IFN-γ, and cytotoxic GrzB, compared to IM-Prime/IN-Boost groups (**fig. S6I and S6J**). For a more comprehensive study of secreted TH1/TH2 cytokine profile, the supernatants of restimulated lung cells were analyzed with multiplexed cytokine assay to assess various cytokine concentrations (**fig. S7, F to K**). Secreted cytokine profile is more associated with TH1 type response with both IM-Prime/IN-Boost, and IN-Prime/IM-Boost groups, which secretes TNF-α and IFN-γ cytokines, respectively. A very low level (or below the detection limit) of TH2 cytokines (IL-4, IL-5, IL-13, IL-10) were observed with IN-Prime/IN-Boost group. These results demonstrate the enhanced and potent TH1 type response in T cells, elicited through IN-Prime/IN- Boost group with PUUC+CpG PAL-NP vaccine formulation.

## DISCUSSION

The continuous waning of pre-existing systemic immunity and immunovasion reduces the current SARS-CoV-2 vaccine’s efficacy for preventing viral transmission. To combat this, vaccines that target the mucosal tissues and induce robust and balanced mucosal-systemic immunity are required to reduce viral shedding, prevent initial infection, and provide overall protection (*7, 8, 17, 52*). Protein subunit nanovaccines formulated with an appropriate combination of adjuvants can represent a promising strategy for developing mucosal vaccines (*53, 54*). Moreover, other specific factors such as nanovaccine design, immunization methods (different prime-boost regimen), respective vaccine doses, and booster time intervals also play a significant role in modulating the SARS-CoV-2 immune response (*54–56*). Therefore in this study, we focus on three important aspects of developing a potent SARS-CoV-2 mucosal subunit nanovaccine: i) designing the nanoparticle by functional group modification in biopolymer that enhance adjuvant delivery and immunomodulation (ii) screening multiple adjuvant combinations of RLRs and TLRs agonists on nanoparticles for enhanced SARS-CoV-2 mucosal immune responses, and (iii) vaccine administration route-specific (prime and boost) comparative study using screened combination adjuvants NPs.

Proper design and synthesis of polymeric biomaterial-based nanocarriers are essential for the enhanced delivery of multiple combination adjuvants that modulate the immune response (*53, 54, 57*). Cationic polysaccharide biomaterials are commonly used for mucosal delivery due to their excellent mucoadhesive property and adjuvanacity (*58, 59*). However, the high number of primary amines in polysaccharides can generate systemic toxicity, which can be reduced by chemical modification with higher-order amines (like secondary/tertiary) (*60*). Therefore, we chose the polysaccharide as a base polymer for NP synthesis and performed the chemical/structural/functional modification to reduce toxicity and enhance multiple adjuvants loading/delivery capability on NPs. This results in an amphiphilic polymer known as polysaccharide-amino acid (arginine/histidine)-lipid polymer (PAL polymer), which undergoes the self-assembly process to form the PAL nanoparticles. Each chemical modification in the polysaccharide has its specific role: (a) the stearyl lipid core provides nanoparticle stability and better encapsulation of hydrophobic adjuvants (TLR7/8 agonist- R848), b) the cationic arginine/histidine aid in surface loading of anionic adjuvant (PUUC RNA: RIG-I agonist, CpG DNA: TLR9 agonist) and provide buffering capacity/endosomal escape for enhanced adjuvant delivery, c) the incorporation of disulfide bond maintains the NP degradability and enhances the adjuvant’s delivery properties. This improved rational design of polymer and nanocarrier will aid in achieving a balanced multiadjuvant delivery, demonstrating minimal toxicity in vitro and in vivo, and ultimately enhancing vaccine effectiveness.

Initial in vitro investigations indicate that adjuvanated PAL-NPs targeting RIG-I and TLRs in different combinations such as: PUUC, R848, PUUC+CpG, and R848+CpG, elicit more diverse proinflammatory cytokine responses (including IL12p70, IL-1β, and IFN-β) with GM-CSF differentiated BMDC, likely due to a broader population of APCs as reported previously for other NP systems such as PLGA or chitosan (*38, 42, 43*). Therefore, we envisioned that these adjuvanated PAL-NP groups would also induce more robust humoral and cellular responses *in vivo*. However, additional studies would be required to determine the efficacy, safety, and potential for their use in future vaccine development. Thus, we designed an in vivo SARS-CoV-2 vaccination study using adjuvanted PAL-NP formulations in a mouse model and conducted two distinct experiments. To conduct in vivo studies, the PAL nanovaccine formulation was prepared by mixing recombinant and stabilized S1 trimer subunit of the SARS-CoV-2 pathogen with multiadjuvanated PAL-NPs. For the physiological resemblance of multiadjuvanated PAL-NPs with the SARS-CoV-2 pathogen (containing ssRNA), a RIG-I agonist PUUC (ssRNA) is selected as one of the major adjuvants. Cytosolic RIG-I-like receptors recognize PUUC RNA and activate them to induce potent antiviral immunity (*38, 39*). In the first in vivo study, we thoroughly investigate the combination adjuvant-mediated immunomodulation using four adjuvanated (R848, PUUC, R848+CpG, PUUC+CpG) PAL nanovaccine formulations, after administrating them in mice with IM-Prime/IN-Boost strategy. This study helps to choose the best multiadjuvant PAL nanovaccine candidate, which strengthens the existing IM immunity and triggers the potent mucosal immune responses against SARS-CoV-2. In the second in vivo study, we conducted a comprehensive head-to-head comparative study of modulated immune responses with different prime-boost immunization routes using the best multiadjuvant nanovaccine candidate examined from the first in vivo study.

Results from the first in vivo study revealed that the PUUC+CpG PAL-NPs is the best multiadjuvanted nanovaccine, as this formulation elicits strong mucosal humoral (IgA and IgG), systemic humoral (IgG and nAb), and local cellular responses. Mucosal IgA is the major antibody response that restricts the entry of respiratory viral pathogens (like influenza, SARS-CoV-2, and MERS CoV) and is the most dominating antibody during early immune response after infection, thus providing sterilizing immunity (*15, 30, 61*). The PUUC+CpG PAL-NPs group also induced TH1 polarized antibody response (IgG2a switching) in both BAL fluid and serum. Prior research has also shown comparable outcomes with adjuvant-based influenza vaccines containing RIG-I agonists that enhance the induction of IgG2a subclass response (*62*). Despite being a preclinical study on mice, our present work can be correlated to human humoral responses, especially the human IgG1, which is analogous to mice IgG2a isotypes and is the most preferred subclass of IgG antibody exhibits optimum antiviral activity (*63*). These findings further highlight the importance of RIG-I agonists in fighting respiratory viral diseases. IN-boosted, PUUC+CpG PAL-NPs after IM priming are the potent inducer of local T cell responses (CD4^+^, CD8^+^, and CD4^+^CD44^+^ TRM) compared to other adjuvant combinations except for PUUC PAL-NPs, which also induces a significant CD4^+^ TRM population. The CD8^+^ tissue-resident memory T (TRM) cells are known to be more effective for viral clearance, and CD4^+^ TRM is involved in a broad spectrum of activities, including the durability of neutralizing antibody responses and promoting the development of protective memory B cells (*49, 64–66*).

Previous studies on some adjuvanated vaccines (alum and CpG adjuvanated) for RSV and SARS-CoV have shown that vaccine-associated enhanced respiratory disease (VAERD) in patients has a connection with CD4 TH2 type response (*67–69*). However, our results show that the RIG-I targeted PUUC+CpG and PUUC PAL-NP group induces TRM, that expresses TH1 type cytokines, and similar results were observed with secreted cytokine expression profile which shows an elevated level of TH1 type cytokine with suppressed or very low detection level of TH2 type cytokines. PUUC+CpG PAL-NPs group also elicits CD8^+^CD44^+^ cells that express higher cytotoxic molecules, like GrzB, and with PUUC PAL-NP group, the induced CD4^+^CD44^+^ cell populations express GrzB. However, CD8^+^ cytotoxic T cells are classically associated with virus-infected cell killing, and CD4^+^ GrzB cytotoxic T cells could be a significant part of the human antiviral T cell responses (*70, 71*). Therefore, both the PUUC+CpG and PUUC PAL-NPs groups, which share a RIG-I agonist as a common adjuvant, significantly enhance the magnitude of TRM responses, polarizing it to TH1 profile, and lead to potent antiviral immunity without showing pathogenic TH2 type responses. Induction of antigen-specific RBD tetramer^+^ B cells with PUUC+CpG PAL-NP group signifies the antigen encounter and further B cell activation and formation of memory B cells (*51*). PUUC+CpG PAL-NP group also enhances the induction of IgM^+^ BRM, IgG^+^ BRM, and IgG^+^ ASC. During reinfection, BRM cells are known to produce rapid and immediate recall responses against pathogen entry at mucosal tissues (*51*).

Although adjuvants can play a crucial role in enhancing potent antiviral mucosal immunity, most studies investigating their effectiveness have focused on IM vaccines with limited knowledge about the role of adjuvants in mucosal vaccines (*31–34, 72*). For example, in humans, CpG (TLR9 agonist) based subunit vaccines elicit a systemic immune response when administered IM and are not an ideal adjuvant candidate for IN immunization (*59*). Few recent studies focused on RIG-I and TLRs targeted SARS-CoV-2 protein-subunit vaccines that can provide a useful comparison point for our study. Nguyen et al. developed a C.S./CpG/RBD intranasal vaccine that induces mucosal humoral response and has demonstrated efficacy against VOCs but lacks to generate local T cell immunity (*35*). Jangra et al. presented a preclinical study showing that a three-dose nanoemulsion RIG-I agonist (IVT DI) adjuvanated SARS-CoV-2 subunit IN vaccine results in systemic TH1 and only IgG responses in both BAL fluid and serum. Despite using multiple high antigen doses of S1 subunit protein and incorporating RIG-I agonists, the study did not observe an enhanced lung-specific humoral and memory T cell response (*36*). The aim of using mucosal vaccines is to establish protective local antiviral immunity. While preclinical studies have shown some success in achieving this, they have not fully met the criteria for overall protection. Therefore, it is essential to enhance current SARS-CoV-2 vaccine strategies by incorporating combination adjuvants.

In contrast, a recent preclinical study by Tianyang et al. showed that IM priming of mRNA LNP and IN-Boost spike protein only (unadjuvanated) elicits protective SARS-CoV-2 immunity (*22*). In our study, we studied how adjuvants, specifically combination adjuvants induce or improve mucosal and systemic immunity. We showed that multiadjuvanated PUUC+CpG PAL- NP group offers an elevated level of mucosal antiviral immunity, both humoral and cellular, as well as a systemic humoral immune response with the IM-Prime/IN-Boost strategy. Also, robust and broad local T cell and comparable mucosal humoral (IgA and nAb) immune responses are induced with IN-Prime/IN-Boost. In comparison to the unadjuvanted approach, our results have revealed significant benefits of using adjuvants. RIG-I is triggered by the native SARS-CoV-2 virus, so we hypothesized that it can mimic infection. Other reported adjuvant formulations, such as BECC: TLR4 agonist and cationic liposome (CAF01), also enhanced mucosal immunity, but with limited local T cell responses (*73, 74*). The recently approved mucosal vaccines by India, China, and others in clinical trials appear to prioritize the induction of a mucosal humoral immune response as indicated by immune markers such as IgA, IgG, and nAbs (*24, 25*). However, the development of robust resident memory T cell responses in the nasal-associated lymphoid tissue (NALT) and lungs provide a more favorable immune signature for potential long-term immunity and better control of lung infections which could lead to stronger protection compared to circulating cellular responses (*16, 22, 47*). Our study also confirmed that the IM-Primed/IN-Boosted PUUC+CpG PAL-NP group induces high-quality, robust mucosal and systemic antibody responses, potent cellular response, and heavily favoring TH1 responses, making it more suitable for a mucosal subunit vaccine.

Different vaccine administration routes have different mechanisms to induce an immune response (*75*). We also confirmed this with our second in vivo study results, where PUUC+CpG PAL nanovaccine formulation was administered in mice via three different prime-boost routes: IM-Prime/IN-Boost, IN-Prime/IN-Boost, IN-Prime/IM-Boost. Our study found that IN-Boosting effectively and significantly enhanced the IM-prime generated immunity (IM-Prime/IN-Boost) with multiadjuvant PUUC+CpG PAL-NP group, which is consistent with our first in vivo study and recent studies available in the literature (*22, 74*). Interestingly, after IN priming, the IN-Boosted mice generated a higher level of lung T cell immunity than IM-Prime/IN-Boost group, along with good local humoral responses (IgA and nAb). The induction of lower T cell response with IN-Prime/IM-Boost vaccination indicates that IM-Boost did not enhance the T cell immunity generated by IN-prime, but rather, the overall response was significantly elevated with IN-Boost as observed in IN-Prime/IN-Boost group. This finding is an interesting point to consider.

Along with a strong T cell response and a comparable mucosal humoral response, the IN- Prime/IN-Boost group shows a considerable systemic IgG response but a relatively lower systemic nAb response than the other two groups, which include IM vaccination in either the prime or boost. The IN-Prime/IN-Boost group enhances the production of monofunctional and polyfunctional subsets of TRM cells (CD4^+^ and CD8^+^) that express TH1 type intracellular cytokines: TNF-α, IFN- γ, and GrzB, but not the pathogenic TH2 type. Their levels are higher than those seen in the IM-Prime/IN-Boost and IN-Prime/IM-Boost groups. A similar trend of T cell and cytokine data was observed in the study on recovered SARS-CoV-2 patients by Grifoni *et al*., which showed that T cell responses appeared as TH1 phenotype with lower levels of TH2 type response (*65, 76*).

Additionally, most IFN-γ^+^CD8^+^ T cells co-expressed GrzB and TNF-α in the same study, which persists in our results also (*65, 76*). The large population of polyfunctional T cells producing multiple TH1 cytokines demonstrates the potential for adequate antigen-presentation and strong co-stimulation from professional APCs (*65*). As SARS-CoV-2 continuously evolves with new immune evasive variants, the population needs to be boosted with the new-generation potent mucosal vaccines that provide enhanced protection and reduced transmission while maintaining their safety and efficacy. Although our results focused on only nanoparticle-based subunit vaccines, if used as a booster, we believe this design would broadly apply to other primary immunization methods. Our preclinical study proves that the mucosal (IN) route is essential for new vaccination strategies and should be included in the current immunization protocols.

In conclusion, multiadjuvanted PUUC+CpG PAL-NP based subunit mucosal vaccine induces robust and potent antiviral mucosal immunity against SARS-CoV-2. We developed an immunization strategy where a balanced mucosal-systemic immunity is induced by the PUUC+CpG nanovaccine when delivering parenteral prime and intranasal boost. Promising outcomes from the intranasal prime and boost nanovaccine delivery also suggest the possibility of a fully mucosal delivery route. These results ensure that this uniquely designed multiadjuvant nanoparticle has great potential to use them for SARS-CoV-2 mucosal subunit nanovaccines. Our results are highly promising in preclinical studies, but due to the inherent immunological differences between animal models and humans, it requires further optimization for future clinical and translational use.

## MATERIALS AND METHODS

All animal experiments were conducted in accordance to approved IACUC (Institutional Animal Care and Use Committee) protocols by the Georgia Institute of Technology.

### Synthesis of PAL-NPs and multiple adjuvant loading

Amphiphilic polysaccharide-amino acid-lipid polymer was synthesized and characterized as described in the supplementary information (see supplementary materials, **fig. S1A**). Cationic polysaccharide-amino acid-lipid nanoparticles (PAL-NPs) are synthesized by probe sonication using an amphiphilic PAL polymer with a final concentration of 0.5 mg/ml. The polymer was first hydrated and dispersed overnight in phosphate buffer saline (PBS, pH 7.2, 10 mM). The hydrated polymer was mixed with DMSO (PBS: DMSO ratio, 80:20), and probe sonicated on ice for 10 min. Nanoparticles were purified with vigorous dialysis in PBS (pH 7.2, 10 mM) for one day by changing water thrice. R848 adjuvant encapsulated cationic PAL-NPs (0.5 µg R848 per mg) were synthesized by the addition of R848 stock in DMSO, followed by probe sonication and dialysis. Nanoparticles were concentrated accordingly to the volume required for the in vivo and in vitro studies. Nanoparticles were electrostatically loaded with nucleic acid adjuvants, either CpG ODN 2395 (Invitrogen, Cat# tlrl-2395) or PUUC (*38, 42*) in 10 mM sodium phosphate buffer (made with nuclease-free water) and left for rotation for 24 h (See table S1 for adjuvant doses). All adjuvants and antigen stock (except R848) were prepared in nuclease-free water. PUUC RNA was synthesized and characterized by following the previously published procedure (*42*). Characterization of adjuvant loading on nanoparticles was described in the supplementary information.

### In vitro activation of mouse BM-APCs with multiadjuvanatyed PAL-NP formulations

GM-CSF-derived BMDCs were generated following previously published procedures (*77*). At day 7 of the culture, mBMDCs derived from GM-CSF were seeded in 96-well plates at a density of 500,000 cells per well and allowed to settle for 2 h. Adjuvanated PAL-NP formulations (see table S1 for adjuvant and PAL-NP doses) were then added to the wells. After treatment, the supernatants were collected 24 hours later, and cytokine concentrations (IL-1β, IFN-β, and IL12p70) were measured using ELISA assays.

### In vivo vaccination studies

BALB/c female mice (6-8 weeks old Jackson Labs, Bar Harbor, ME) were used to measure the adaptive immune response and anesthetized using 30% v/v Isoflurane diluted in propylene glycol for vaccination. Final nanovaccine formulations are prepared by mixing the adjuvanated PAL-NPs (250 µg per mice) and recombinant, stabilized SARS-CoV-2 spike S1 trimer protein (1µg/mice, SPN-C52H9, ACRO Biosystems). For intramuscular vaccination, formulations were prepared in total 100 µl of PBS (pH 7.2, 10 mM), out of which 50 µl was injected to the right and 50 µl to the left anterior tibialis muscle at day 0 as the first dose and at day 21 for boost doses). The doses of adjuvants on the PAL-NP adjuvants formulation for the IM vaccination (per mice) are PUUC (20 µg), CpG (40 µg), and R848 (20 µg), and also shown in **table S1**. For intranasal vaccination, the formulations are prepared in a total of 40 µl of PBS (pH 7.2, 10 mM), out of which 20 µl was administered dropwise in both the left and right nares. The doses of adjuvants on the PAL-NP adjuvants formulation for the IN vaccination (per mice) are PUUC (20 µg), CpG (20 µg), R848 (20 µg), as shown in table S1.

### Ex vivo lung cell restimulation and T cell staining

After harvesting the lungs from vaccinated mice (individual experiment), single-cell suspensions were prepared with a gentleMACS™ Octo Dissociator and Lung Dissociation Kit (Miltenyi Biotec) according to the manufacturer’s instructions, including RBC lysis. Cells were centrifuged and resuspended at 10 million cells/mL in RPMI media with 10% FBS, 1% penicillin-streptomycin, 1 mM sodium pyruvate, and 1x β-mercaptoethanol. Cells were seeded at 2 million cells per well in a U-bottom 96-well plate and left to culture overnight (at 37^◦^C with 5% CO2).

Lung cells were centrifuged and resuspended with fresh complete RPMI media with 20 μL/mL of PeptTivator® SARS-CoV-2 Prot_S (Miltenyi Biotec) and 5 μg/mL Brefeldin A (Biolegend). After incubation for 6 h, cells were stained for 30 min at RT with Zombie Green™ Fixable Viability Kit (Biolegend) and were blocked with anti-mouse CD16/32 (Biolegend) and True-Stain Monocyte Blocker™ (Biolegend). For blocking of cell surfaces, the cells were stained for 30 min at 4°C with surface antibodies: anti-mouse CD3 (Biolegend, APC Fire 810), CD4 (Biolegend, APC), CD8a (Biolegend, PE/Cy5), CD44 (Biolegend, BV711), CD69 (Biolegend, BV785), CD103 (Biolegend, PE-Dazzle 594), CD56 (BD, BUV395), and TCR-γδ (Biolegend BV510). After surface staining, the cells were stained for intracellular cytokines. The cells were fixed and permeabilized for 30 min with the Foxp3/Transcription Factor Staining Buffer Set (eBioscience) at 4°C. Then cells were stained with anti-mouse TNF-α (Biolegend, PE/Cy7), IFN-γ (Biolegend, PE), and Granzyme B (Biolegend, Pacific Blue). Cell population data were acquired on Cytek Aurora flow cytometer and analyzed using FlowJo Software (**fig. S8A** for the gating strategy).

### B cell staining and flow cytometry

Lung single-cell suspensions were stained for 30 min at RT with Zombie Red™ Fixable Viability Kit (BioLegend) and were blocked with anti-mouse CD16/32 (Biolegend) and True-Stain Monocyte Blocker™ (Biolegend). Cells were washed once with PBS before surface staining. Following blocking, cell surfaces were stained for 30 min at 4°C with anti-mouse GL7 (Biolegend, Pacific Blue), IgM (Biolegend, Pe-Cy7), CD138 (Biolegend, PerCP/Cy5.5), CD19 (Biolegend, AF700), IgA (SouthernBiotech, FITC), B220 (Biolegend, BV711), CD38 (Biolegend, APC Fire 750), and anti-IgD (Biolegend, BV605), PE-SARS-CoV-2 RBD tetramer, APC-SARS-CoV-2 RBD tetramer for 30 min at 4°C (tetramer synthesis is described in supplementary information). After washing with PBS, cells were fixed using 4% paraformaldehyde. Cell population data were acquired on Cytek Aurora flow cytometer and analyzed using FlowJo Software (**fig. S8B** for gating strategy).

### ELISA assay for quantifying anti-spike antibody responses

The diluted recombinant SARS-CoV-2 Spike His Protein, CF (R&D Systems, Cat# 11058-CV) (1 μg/mL in 0.05 M carbonate-bicarbonate buffer, pH 9.6) was coated onto Nunc™ MaxiSorp™ ELISA plates by adding 100 ng/well and incubating the plates overnight at 4°C. Antigen-coated plates were washed three times with PBST wash buffer (prepared by mixing 10 mM PBS and 0.05% Tween-20), and plates were blocked for six hours at 4°C with PBSTBA (prepared by mixing PBST with 1% BSA and 0.02% NaN3). Blocked plates were incubated overnight at 4°C with diluted serum and BAL fluid samples (individual experiments). Plates were washed three times with PBST. A secondary biotinylated anti-mouse IgA, total IgG, IgG1, or IgG2a antibody (SouthernBiotech) which is 5,000-fold diluted in 5-fold diluted PBSTBA, was added to plates for 2 h at RT. Plates were similarly washed with PBST. After two hours, a 5,000- fold diluted streptavidin-conjugated horseradish peroxidase (strep-HRP, ThermoFisher) was added to the plates and incubated for the next 2 h at RT. The plates were again washed six times. Ultra TMB-ELISA Substrate Solution (ThermoFisher) was incubated for 15 to 20 min for color development on the plate. Lastly, stop solution (2 N sulfuric acid) was added to each well, and absorbance was measured at 450 and 630 nm (background) on a Synergy H.T. plate reader (BioTek) with Gen5 software.

### Modified ELISA assay to measure anti-spike neutralizing antibodies

For the quantification of neutralizing antibodies, the above-described ELISA method is used with slight modification. A diluted recombinant SARS-CoV-2 Spike His Protein, CF (R&D Systems, Cat# 11058-CV) in 0.05 M carbonate-bicarbonate buffer (1 μg spike/mL, pH 9.6) was incubated in wells of a Nunc™ MaxiSorp™ ELISA 96-well plate (100 ng/well) overnight at 4°C. Plates were washed three times with PBS-Tween wash buffer (PBST). Antigen-coated plates were blocked in PBSTBA for six hours at 4°C. Blocked plates were again incubated overnight with serum and BAL fluid samples (individual experiments). Plates were similarly washed. Plates were incubated with PBSTBA diluted 500 ng/mL (25 ng/well) recombinant biotinylated human ACE-2 (R&D Systems, Cat# BT933-020) for 2 h at RT. The plates were washed again with PBST. A 5,000-fold diluted strep-HRP (ThermoFisher) was added to the plates and incubated for 2 h at RT. Plates were extensively washed six times and incubated with the optimized volume of Ultra TMB-ELISA Substrate Solution (ThermoFisher) for 20 min. In the end, the reaction was stopped with a stopping reagent (2 N sulfuric acid), and absorbance was measured at 450 and 630 nm (background) on a Synergy H.T. plate reader (BioTek) with Gen5 software.

### Statistical analysis

All flow cytometry FCS files were analyzed with FlowJo (v10, B.D.). Statistical analyses were performed with GraphPad Prism 9. For comparisons of more than two groups, statistical differences between normally distributed datasets were determined with a one-way analysis of variance (ANOVA) followed by Tukey’s and Bonferroni’s post hoc test for multiple comparisons. Similarly, nonparametric datasets were evaluated with the Kruskal-Wallis test and Dunn’s post-hoc test. *P* < 0.05 was considered statistically significant.

### Data and materials availability

All data needed to evaluate the conclusions in the paper are present in the paper and/or the Supplementary Materials.

## Acknowledgments

We thank Mark Cole Keenum, Alexandra Atalis, and Pallab Pradhan for their advice and consultation during the experiment design and early data analysis. We acknowledge the core facilities at the Parker H. Petit Institute for Bioengineering and Bioscience at the Georgia Institute of Technology for using their services, shared equipment, and expertise. These facilities include the Biopolymer Characterization Core, the Engineered Biosystems Building, Physiological Research Laboratory for animal experiments, and the Cellular Analysis and Cytometry Core for flow cytometry experiments. We also acknowledge the NMR facility at the Molecular Sciences and Engineering (“MoSE”) building at Georgia Institute of Technology. We acknowledge Dr. Leslie Gelbaum and Dr. Johannes Leisen for NMR training, advice, and help with NMR sample preparation and polymer characterization. Structural representations of polymers were drawn in ChemDraw software. Graphpad plots were arranged in Adobe Illustrator, and vaccination figures were made with Biorender.

## Author contributions

Conceptualization and designing: B.P. and K.R. Methodology: B.P. and K.R. Investigation: B.P., Z.W., A.J., E.B., R.J., A.B., D.M., J.H., C.V., L.O.F., D.J.H., R.K.N., R.R., C.S., and M.A.O. Visualization: B.P. and Z.W. Supervision: B.P. and K.R. Writing original draft: B.P. and K.R. Writing, review & editing: B.P. and K.R.

