## Supplementary Materials for "A multiadjuvant polysaccharide-amino acid-lipid (PAL) subunit nanovaccine generates robust systemic and lung-specific mucosal immune responses against SARS-CoV-2 in mice"

*Bhawana Pandey et al.*

**This PDF file includes:**

Supplementary Materials and Methods
Table S1
Figs. S1 to S13
References

### MATERIALS AND METHOD:

#### 1. Materials:

Chitosan polysaccharide (Mw 15KDa) was purchased from Polysciences (85% degree of deacetylation). Dialysis tubing (MWCO 3.5kDa, 10kDa) was purchased from Thermo-Fisher Scientific. NMR solvents and other solvents for synthesis, such as ethanol and diethyl ether, were purchased from Sigma Aldrich.

#### 2. Methods: Experimental section

##### 2.1. Synthetic steps for polysaccharide (chitosan)-amino acid-lipid polymer: multistep synthesis

###### 2.1.1. Synthesis of O-carboxymethyl-chitosan: OCMC

O-Carboxymethyl-Chitosan (OCMC) was synthesized by following the previously described procedure to increase the selective O-carboxylation and reduce the N-carboxylation *(78)*. The C-6 position of chitosan polysaccharide (500 mg) was first alkalized with 50% aqueous NaOH (20 mL) at -10°C for one hour. The alkalized polysaccharide was further reacted with 2.5 g monochloroacetic acid (Sigma Aldrich) at 45-55°C for six h. The reaction mixture was added with 70% ethanol to prepare the sodium salt of OCMC, which was further purified by vacuum filtration. The OCMC sodium salt was washed with 70% ethanol and acidified with 1 N HCl to form OCMC. The obtained OCMC was filtered and dried under a vacuum for further use. The incorporation of O-carboxymethyl group at the C-6 position was confirmed by $^{1}$HNMR (**fig. S9**).

###### 2.1.2. Synthesis of thiolated O-carboxymethyl-chitosan: OCMC-SH

Thiolated OCMC was synthesized by modifying the previously published procedure *(79)*. Briefly, synthesized OCMC was thiolated by covalent conjugation of carboxyl of thioglycolic acid with the amine group of chitosan (C-2 position) using carbodiimide chemistry. Firstly, the carboxyl group of TGA (500 mg, Sigma Aldrich) groups was activated with 1-ethyl-3-(3-dimethylaminopropyl) carbodiimide hydrochloride (EDC, Thermo Fischer) at pH 6.5 in DI water with a final concentration of 125 mM for 2 h. 250 mg of OCMC was acidified with 1M HCl. OCMC solution was added to the activated TGA solution, and the pH of the reaction medium was adjusted to 5 to avoid forming the disulfide bond. To eliminate excess TGA and to purify the thiolated OCMC, the reaction mixtures were dialyzed five times using dialysis membrane 10 kDa MWCO (Sigma Aldrich) for two days in the dark against HCl (5 mM), then two times against HCl (5 mM) with 1% NaCl at 10°C, which helps quench the ionic interactions between anionic sulfhydryl and the cationic polymer. Final dialysis was performed against 1 mM HCl to maintain the pH of the thiolated OCMC polymer to 4. Polymers were further lyophilized and stored at 4°C until further use. Thiolation was confirmed by using $^{1}$HNMR (**fig. S10**) and Elmann's assay (**fig. S1B**).

###### 2.1.3. Synthesis of cysteamine conjugated OCMC: OCMC-S-S-Cys

The above lyophilized thiolated OCMC was first reduced with DTT (Dithiothreitol, Sigma Aldrich) before cysteamine conjugation. The necessary reduction step reduces the disulfide bond formed during lyophilization and helps increment free sulfhydryl groups. For reduction, the thiolated OCMC-SH solution was prepared in DI water, and pH was maintained at 8 using 1 M NaOH. DTT (Dithiothreitol, Sigma Aldrich) was added in a final concentration of 100 mM, and the reaction mixture was continuously stirred at RT. After approximately two h, NaCl was added to the reaction mixture (final concentration 1%, weight/volume), and the pH was adjusted to 4 with 1 M HCl. The resulting solution was dialyzed using a similar method as discussed in step 2.1.2. Finally, the solution was filtered through a 0.5 $\mu$m filter. The second cycle with DTT was repeated to improve the reduction of remaining disulfide bonds. The filtered thiolated OCMC was conjugated to cysteamine by forming a disulfide bond between thiols of cysteamine (Sigma Aldrich) and free thiols on OCMC. Cysteamine solution was prepared in 1% acetic acid, and the solution pH was adjusted to 6.0. The thiolated OCMC was added dropwise over a three h period using an addition funnel to the cysteamine solution. The reaction mixture was allowed to stir for 24 h at RT, and the pH of the final solution was maintained between 4-5. The resulting polymer conjugates were isolated using a similar dialyzing method, as discussed in step 2.1.2. Polymers were further lyophilized and stored at 4°C until further use. Synthesis of OCMC-S-S-Cys was confirmed by using $^1$HNMR (**fig. S11**) and Elmann's assay (**fig. S1B**). The concentration of thiols was decreased after disulfide formation, as shown by Elmann's assay, which confirms the synthesis of OCMC-S-S-Cys formation.

**2.1.4. Synthesis of arginine and histidine conjugated OCMC-S-S-Cysteamine: OCMC-S-S-(A/H)**

The coupling of both amino acids: N-Boc Histidine (Alfa Aesar) and N-Boc Arginine (Alfa Aesar), onto cysteamine conjugated OCMC was performed by the reaction of the amine groups of OCMC-S-S-Cys and carboxylic group of amino acid in the presence of coupling agents EDC (Thermo Fischer Scientific) and NHS (Sigma Aldrich). The Boc-protected amino acids were used to reduce the cross-reaction of the free carboxyl group of the C-6 position of chitosan with the amine groups of amino acids. The free carboxyl group of N-Boc Histidine (0, 5, 10 mM) and N-Boc Arginine (0, 10, 20 mM) was first activated individually by the addition of EDC/NHS (10 molar excess) in TEMED/HCL buffer (1% concentration, v/v) at pH 5.5 (Tetramethylethylenediamine, Sigma Aldrich) for 2 h at 25°C. The activated amino acid solution was added dropwise to the solution of OCMC-S-S-Cys in the same buffer and reacted for the next 16 h. The concentration of both amino acids was used with different ratios to yield conjugates with different degrees of substitution. The pH of the final reaction mixture was maintained at 6. Then the reaction product was extensively dialyzed against distilled water for two days, and the pH was maintained at 6.5 to remove the unreacted components. The purified polymer was then recovered by lyophilization and stored at 4°C. The incorporation of arginine and histidine in the polymer chain was confirmed by $^1$H NMR (**fig. S12**).

**2.1.5. Synthesis of polysaccharide-amino acid-lipid (PAL) polymer: OCMC-S-S-(A/H)-SA**

Chit-S-S-(A/H)-SA was synthesized by the coupling of the carboxyl group of OCMC-S-S-(A/H) with the amine group of stearyl amine (TCI Chemicals). In brief, the carboxyl group of OCMC-S-S-(A/H) (250 mg) at the C-6 position was activated by the addition of EDC and NHS in 20 mL PBS (pH 6, 10 mM) for two hours. Different amount of stearyl amine (0.25–0.625 mol/mol glucosamine residues) was used to react with -COOH groups of OCMC-S-S-(A/H). The stearyl amine was pre-dissolved in 20 mL ethanol by heating at 60°C in a separate round bottom flask. After two hours, the stearyl amine solution was added dropwise to the OCMC-S-S-(A/H) polymer solution by maintaining a similar temperature at 60°C and again heated to 80°C for the next 6 h. After that, the reaction mixture was allowed to cool to room temperature and again stirred for 18 h. For purification, the reaction mixture was vigorously dialyzed (MWCO 3.5 KDa) against distilled water for 48 h to remove water-soluble by-products and ethanol. The dialyzed suspension was lyophilized and rinsed several times with hot ethanol and diethyl ether and precipitated in ethanol to remove unreacted stearyl amine. Boc deprotection of the amino acids conjugated at the C-2 position of OCMC-S-S-(A/H)-SA (200 mg) was performed using 2 M HCl in dioxane (2 mL) and trifluoroacetic acid (TFA) in ice-cold temperature under an argon atmosphere and stirred for 15 min and further stirred for next three hours at RT. The reaction product was further precipitated in ethanol, washed, and dried. The residue was dialyzed against 0.01 N HCl by redissolving in DI water using dialysis tubing of 3.5 kDa MWCO. The samples were initially dialyzed against 0.01 N HCl for one day and then with DI water for another day with several water changes. $^{1}$H NMR confirmed the incorporation of stearyl chains in the OCMC-S-S-(A/H) polymer (**fig. S13**).

### 2.2. Characterization Methods

#### 2.2.1. Quantification of free thiol content in polymers: Elmann's assay

To confirm the synthesis of OCMC-S-S-Cys and reduced concentration of free thiols in OCMC-S-S-Cys polymer compared to OCMC-SH, we performed the Elmann's assay of OCMC after and before thiolation as well as after disulfide bond formation. In brief, a reaction buffer was prepared using sodium phosphate 0.1 M and 1 mM ethylenediaminetetraacetic acid (EDTA) in pH 8.0. The stock solution of Elmann's reagent was prepared by dissolving 2 mg of 5,5′-dithiobis-(2-nitrobenzoic acid) in 0.5 mL same reaction buffer. Per the manufacturer's instructions, 500 µL of the sample (1mg of polymer in 1 mL of reaction buffer) was added to a test tube containing 100 µL Elmann's reagent and 5 mL of reaction buffer. The samples were incubated for an optimized time at 37°C and protected from light. A 100 µl of the sample was transferred to a 96-well plate. Samples were analyzed using a microplate reader (BIOTEK Synergy HT plate reader, Gen5 software) at a wavelength of 485 nm to determine the content of thiol groups. For the estimation of disulfide contents, in OCMC-S-S-Cys polymers were first reduced with $NaBH_4$ and then evaluated by Elmann's reagent. A serial dilution of cysteine hydrochloride monohydrate was used as a standard, and a standard curve is generated using eight serial concentrations of 1.5, 1.25, 1.0, 0.75, 0.5, 0.25, 0.125, 0.0625, and 0 mM. All experiments were performed in triplicate. The free thiol content was quantified according to the following equation:

$$Thiolation\ \% = \frac{(OCMC \cdots SH - OCMC)}{(OCMC \cdots SH)} \times 100$$

Where OCMC and OCMC-SH stands for carboxylated chitosan and thiolated carboxylated chitosan, respectively.

#### 2.2.2. Nanoparticles stability

A time-dependent PAL-NPs degradability behavior was evaluated using DTT as a reducing agent. The PAL-NPs (0.5 mg/mL) dispersion with and without disulfide bond was prepared in PBS (10 mM, pH 7.4). The reducing agent dithiothreitol was added to the solution with the final concentration of 10 mM. The samples were incubated at 37°C and protected from light. At regular time points (0 h, 2 h, 6 h, and 12 h), the particle's average size was measured by DLS (Dynamic Light Scattering). Particle size degradation with respect to the time was plotted. (See **fig S1c**).

#### 2.2.3. Adjuvant loading on PAL-NPs

Nanoparticle size and surface zeta potential before the anionic adjuvant loading were measured with a Zetasizer Nano Z.S. (Malvern), as shown in table S1. The sample preparation details for TEM are provided in section 2.2.5. R848 encapsulation was determined by dissolving PAL-NPs particles in DMSO (Tocris, Cat# 3176), followed by absorbance readings against a R848 standard curve at 324 nm. PUUC RNA loading was quantified by Ribogreen assay according to the manufacturer's instructions. CpG DNA loading was quantified by measurement of unbound DNA in the supernatant after centrifugation at 20,000*g*, using a Nucleic Acid Quantification workflow on a Synergy H.T. plate reader (BioTek) with Gen5 software.

#### 2.2.4. NMR spectroscopy

The $^{1}H$ NMR analysis was performed on a Bruker Avance III 400 at 25°C. OCMC, OCMC-SH, OCMC-S-S-Cys polymers were dissolved in $D_2O$ with 1% DCl. OCMC-S-S-(A/H) and OCMC-S-S-(A/H)-SA polymers were dissolved in deuterated dimethyl sulfoxide (DMSO-d6). Chemical shifts were recorded in parts per million (ppm) using the signal of TMS as the internal reference. NMR spectral data were analyzed using MestreNova NMR software.

#### 2.2.5. Transmission electronic microscopy (TEM)

TEM was performed on a FEI Tecnai G2 F30 S-TWIN Transmission Electron Microscope at 300 kV. The 10 µL PAL-NPs solution (10 times a diluted sample of 0.5 mg/mL) was placed on the copper grids for sample preparation. The excess solution was absorbed by Whatman filter paper at the edges and dried for 10 sec at RT. Samples were further stained by a drop of phosphotungstic acid (stock solution of 2%) onto the surface of the sample-loaded grid. The grid was washed twice with DI water to remove excess staining reagent. Grids were dried in desiccators overnight and analyzed by transmission electron microscopy.

### 2.3. Immunological studies:

#### 2.3.1. Euthanasia and sample collection (BAL fluid, blood, and lungs)

For IM and IN in vivo studies, mice were euthanized at day 35 (after 2 weeks of booster dose), and blood, BAL fluid, and lungs were harvested. Mice were initially anesthetized using an optimized mixture of ketamine (80 mg/kg) and xylazine (15 mg/kg), injected 25 ul intraperitoneally first and 50 ul intramuscularly later (7-8 minutes later). Blood was first collected from all mice via the jugular veins. All blood samples were allowed to clot for 30–60 min at RT in serum separator tubes (B.D., #365967), and serum was separated by centrifugation at 4000*g* for 15 min at 4°C. Serum samples were heat inactivated at 56°C for 30 min in a water bath which inhibits the complement binding. After inactivation, serum samples were aliquoted and stored at -80°C. BAL fluid was collected after two separate injections and withdrawals (total 2 ml in Hanks' Balanced Salt Solution, sigma Aldrich cat#H4641 with 100 µM EDTA Sigma Aldrich cat#03699,) by inserting a 20 gauze one-inch catheter into the trachea following the methods described in Hoecke et al *(80)*. Samples were centrifuged at 300g for 5 minutes to remove cells. BAL Samples were further concentrated 10x using 100KDa Amicon concentrators and aliquoted and stored at -80°C.

#### 2.3.2. SARS-CoV-2 RBD B-cell Tetramer Production

RBD tetramer was prepared by a previously published procedure *(22)*. Recombinant Biotinylated SARS-CoV-2 S protein RBD, His, Avitag™ (ACRO Biosystems SPD-C82E9) was incubated at a 4:1 molar ratio with either streptavidin-PE (Biolegend, 405204) or streptavidin- APC (Biolegend, 405207) in PEB buffer (1X PBS + 0.5% BSA 2 mM EDTA) for one hour at 4°C. The mixture was then purified, concentrated in an Amicon Ultra (50 kDa

MWCO) spin column, and washed with sterile, cold PBS. Excess streptavidin was blocked with biotin. Final protein concentration was measured on a nanodrop, using a protein quantification workflow on a Synergy H.T. plate reader (BioTek) with Gen5 software. Tetramers were diluted to 1.0 µM in PBS and stored at 4°C.

#### 2.3.3. Supernatant cytokine profile

Supernatant from the lung T cell restimulation assay were harvested, and $T_H1/T_H2$ cytokine production was measured using LEGENDplex™ (Mouse $T_H1/T_H2$ Panel, Biolegend, 741054) for IL-5, IL-13, IL-2, IL-6, IL-10, IFN-γ, TNF-α, IL-4, according to manufacturer's instructions. Cytokine beads were analyzed on a cytoflex flow cytometer. Raw data were analyzed using LegendPlex software (Biolegend), and the average cytokine level was determined from two duplicate samples.

**Table S1. Adjuvanated PAL-NPs formulations for in vitro and in vivo studies.**

| PAL-NP Formulations | Size (nm)# | Polydispersity Index (PDI) # | Zeta potential (mV)# | Adjuvant dose for BMDCs (ng/5e5 cells/mL) | Adjuvants loading levels for in-vivo studies (ug/mg NPs) |
|---|---|---|---|---|---|
| Blank PAL-NPs | 249.4 ± 31.1 (n=3) | 0.21 | 31.7 ± 2.5 (n=3) | 12ug NPs | 250ug NPs (IN and IM) |
| R848 PAL-NPs | 236.0 ± 65.2 (n=3) | 0.175 | 30.0 ± 6.92 (n=3) | 20ng R848 | 80ug R848/mg NPs (IN and IM) |
| CpG PAL-NPs | 249.4 ± 31.1 (n=3) | 0.21 | 31.7 ± 2.5 (n=3) | 100ng CpG | 160ug CpG/mg NPs- IM<br>80ug CpG/mg NPs- IN |
| PUUC PAL-NPs | 249.4 ± 31.1 (n=3) | 0.21 | 31.7 ± 2.5 (n=3) | 100ng PUUC | 80ug PUUC/mg NPs (IN and IM) |
| R848+CpG PAL-NPs | 236.0 ± 65.2 (n=3) | 0.175 | 30.0 ± 6.92 (n=3) | 20ng R848 and 100ng CpG | 80ug R848 and 80ug PUUC/mg NPs (IN and IM) |
| PUUC+CpG PAL-NPs | 249.4 ± 31.1 (n=3) | 0.21 | 31.7 ± 2.5 (n=3) | 100ng CpG and 100ng PUUC | 160ug CpG and 80ug PUUC/mg NPs- IM<br>80ug CpG and 80ug PUUC/mg NPs- IN |
| R848+PUUC PAL-NPs | 236.0 ± 65.2 (n=3) | 0.175 | 30.0 ± 6.92 (n=3) | 20ng R848 and 100ng PUUC | - |
| R848+CpG+PUUC PAL-NPs | 236.0 ± 65.2 (n=3) | 0.175 | 30.0 ± 6.92 (n=3) | 20ng R848, 100ng CpG and, 100ng PUUC | - |

#Size, PDI, and zeta potential measurements were taken for all NPs prior to electrostatically loading adjuvants PUUC or CpG.

### FIGURES

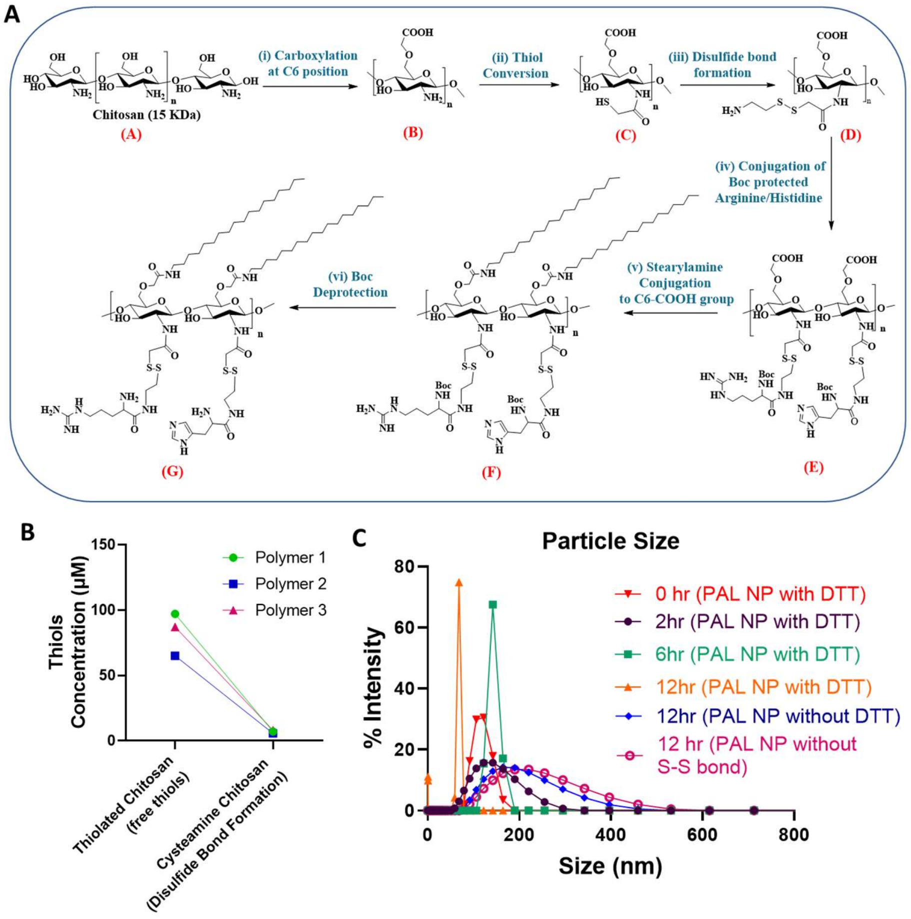


**Figure S1**. **Synthetic scheme of PAL polymer and PAL-NPs characterization.** (**A**) Multistep synthesis of polysaccharide-amino acid-lipid amphiphilic (PAL) polymer- (i) Chitosan, NaOH, -10ºC incubation, 1 h, Cl-$CH_2$-COOH, heat (45ºC), 24 h (ii) EDC/NHS, HS-$CH_2$COOH (iii) $NH_2$-$CH_2$-$CH_2$-SH, cysteamine, pH=6 (iv) EDC/NHS, Nα-Boc-L-arginine and Nα-Boc-L-histidine (v) EDC/NHS, $CH_3(CH_2)_{17}NH_2$, heating 80ºC (vi) TFA/4M HCl in Dioxane, Boc deprotection. (**B**) Estimation of thiols and disulfide concentration in thiolated polymer and cysteamine conjugated chitosan polymer by Elmann assay. (**C**) Time-dependent degradation study of the PAL-NPs by DLS analysis in the presence of DTT (10 mM).

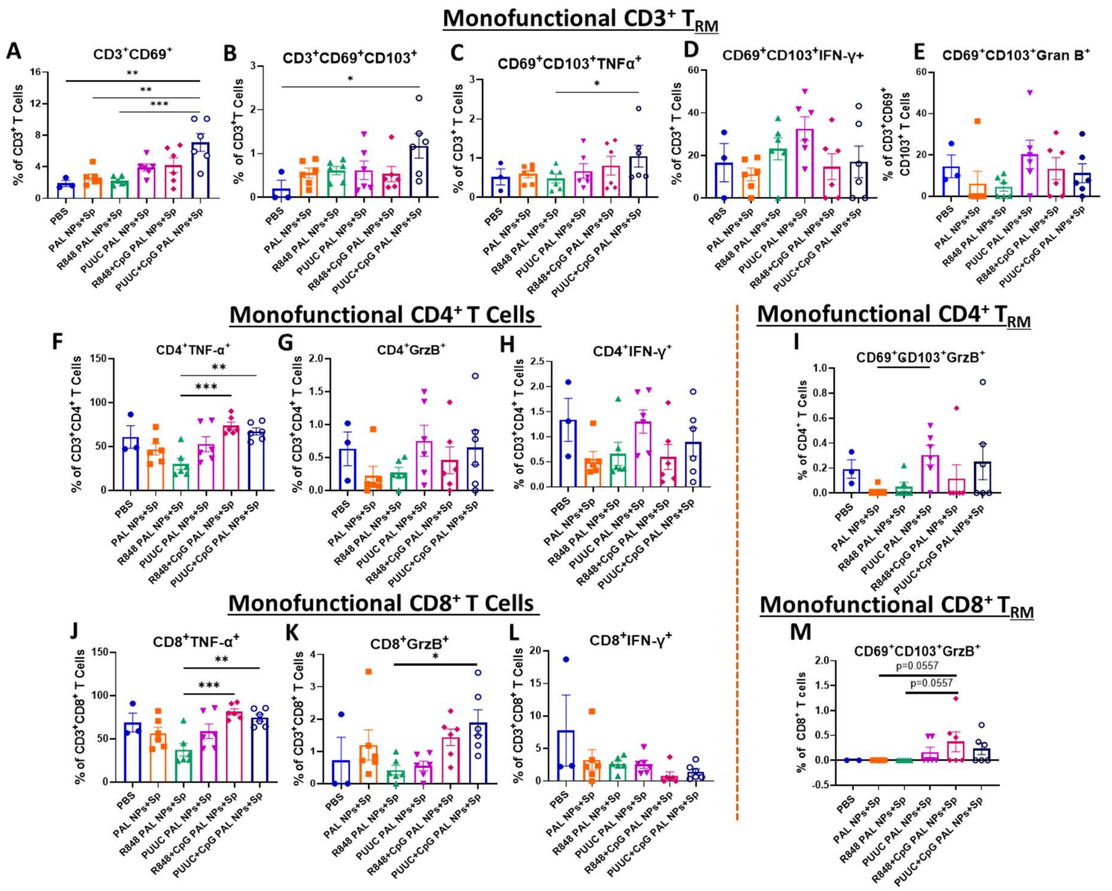


**Figure S2. PUUC+CpG PAL-NP protein subunit vaccine formulation with S1 spike protein, elicits robust SARS-CoV-2 elicits T cell immunity when delivered IM-Prime/IN-Boost**. On days 0 (IM prime) and 21 (IN boost), female BALB/c mice were immunized with adjuvanated PAL-NP vaccine formulation with S1 spike protein (see Materials/Methods and Table 1 for doses). Mice were euthanized, and lungs were collected on Day 35 (one-week post-boost). Lung cells were restimulated with spike peptide for 6 h. (**A** and **B**) Percentage of $CD3^+CD69^+$ and $CD3^+CD69^+CD103^+$ ($CD3^+$ $T_{RM}$) cell population. (**C** to **E)** Percentage of monofunctional $CD3^+$ $T_{RM}$ cells expressing TNF-α, IFN-γ, and GrzB. (**F** to **H**) Percentage of monofunctional $CD4^+$ T cells expressing TNF-α, IFN-γ, and GrzB. (**I**) Percentage of Monofunctional $CD4^+$ $T_{RM}$ cells expressing GrzB. (**J** to **L)** Percentage of monofunctional $CD8^+$ T cells expressing TNF-α, IFN-γ, and GrzB. (**M**) Percentage of polyfunctional $CD8^+$ $T_{RM}$ cells expressing GrzB. Error bars represent the SEM. Statistical significance was calculated using one-way ANOVA followed by Tukey's post-hoc test for the figures (**A**), (**B**), (**J**), and (**K**), and Bonferroni's post-hoc test for the figures (**C**), (**I**), and (**M**), for multiple comparisons. $*p \leq 0.05$, $**p \leq 0.01$, $***p \leq 0.001$, $****p \leq 0.0001$ for all graphs.

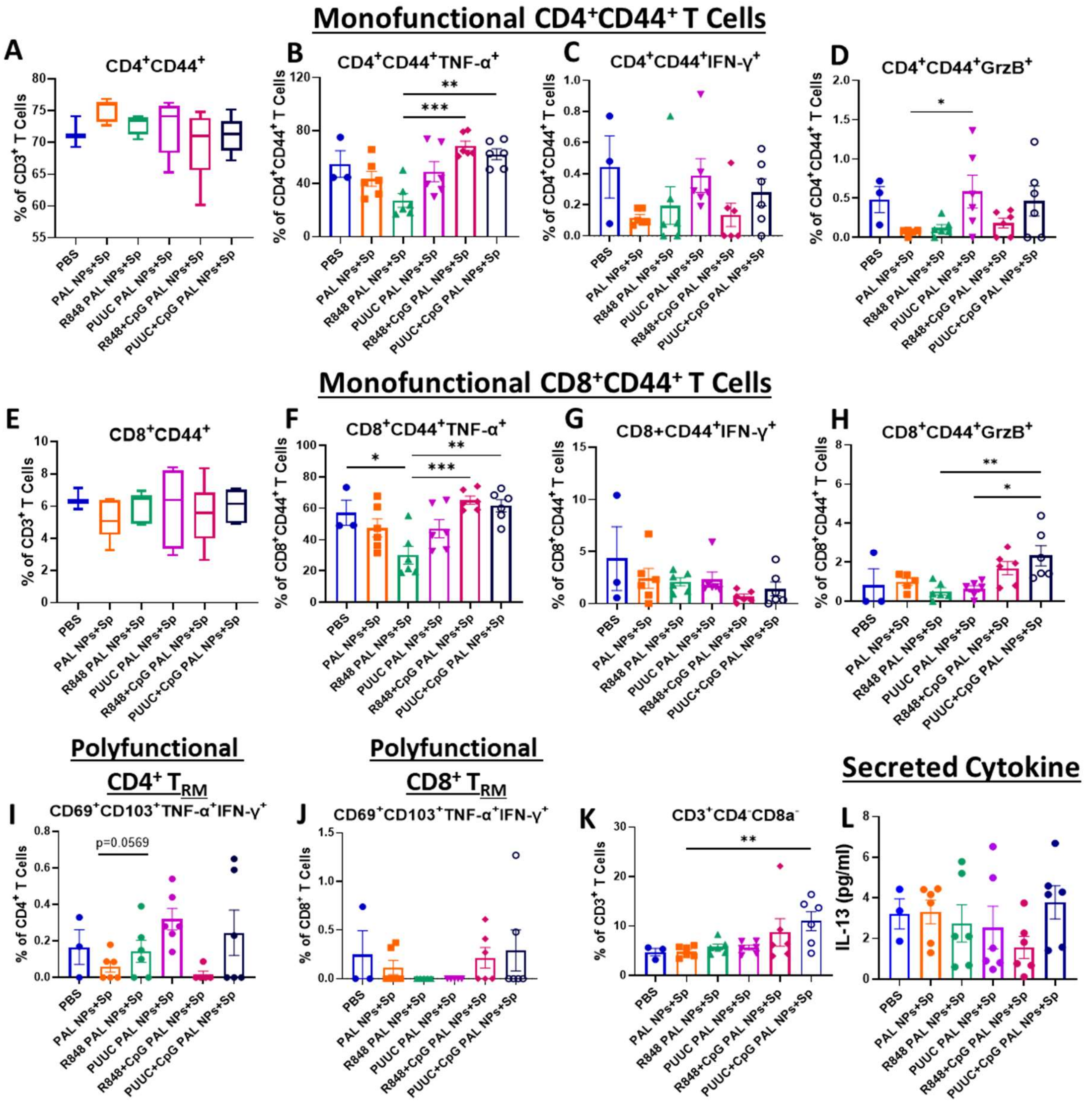


**Figure S3. PUUC+CpG PAL-NP protein subunit vaccine formulation with S1 spike protein, elicits robust SARS-CoV-2 elicits T cell immunity when delivered IM-Prime/IN-Boost.** On days 0 (IM prime) and 21 (IN boost), female BALB/c mice (n=3 for PBS and n=6 for other adjuvanated PAL-NP groups) were immunized with adjuvanated PAL-NP vaccine formulation with S1 spike protein (see Materials/Methods and Table 1 for doses). Mice were euthanized, and lungs were collected on Day 35 (one-week post-boost). Lung cells were restimulated with spike peptide for 6 h. (**A**) Percentage of $CD4^+CD44^+$ cell population. (**B** to **D)** Percentage of $CD4^+CD44^+$ cells expressing TNF-α, IFN-γ, and GrzB. (**E)** Percentage of monofunctional cells expressing $CD8^+CD44^+$. (**F** to **H**) Percentage of monofunctional $CD8^+CD44^+$ T cells expressing TNF-α, IFN-γ, and GrzB. (**I**) Percentages of monofunctional $CD4^+$ $T_{RM}$ cell population co-expressing both TNF-α and IFN-γ. (**J**) Percentage of monofunctional $CD8^+$ $T_{RM}$ cell population co-expressing both TNF-α and IFN-γ. (**K**) Percentage of $CD3^+CD4^-CD8^-$ cell population. (**L**) IL-13 Cytokine concentration in supernatants from restimulated lung cells. Error bars represent the SEM. Statistical significance was calculated using one-way ANOVA followed by Tukey's post-hoc test for the figures (**B**), (**F**), and (**H**), and Bonferroni's post-hoc test for the figures (**D**), (**I**), and (**K**), for multiple comparisons. Statistical significance for cytokine concentrations was calculated with One-Way ANOVA and Tukey post-hoc test. *$p \leq 0.05$, **$p \leq 0.01$, ***$p \leq 0.001$, ****$p \leq 0.0001$ for all graphs.

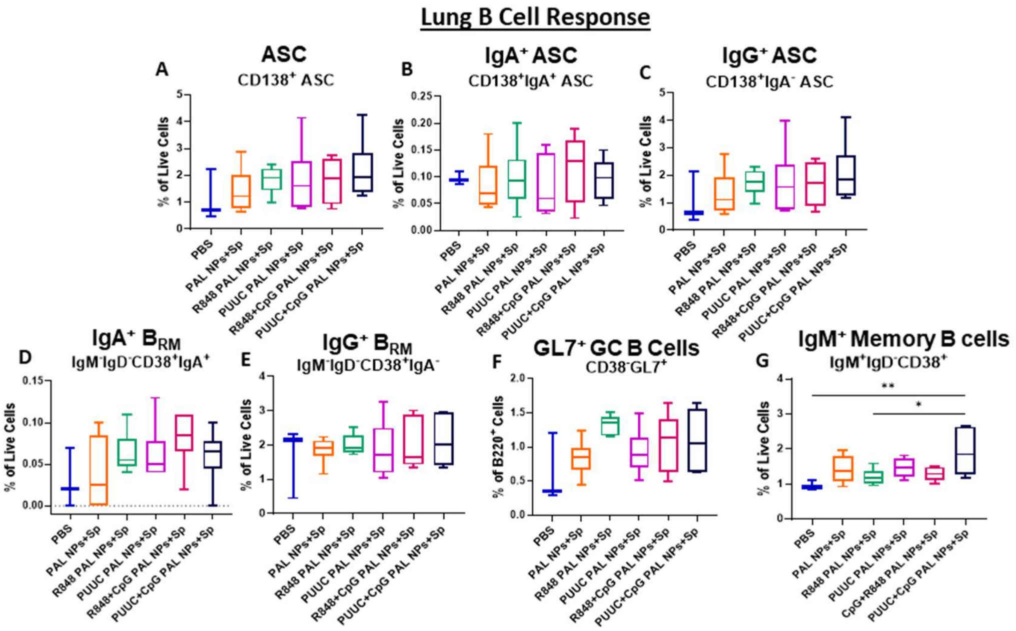


**Figure S4. Analysis of lung B cell responses when multiple adjuvanated PAL-NP protein subunit vaccine formulations are delivered to mice via IM-Prime/IN-Boost vaccination.** On days 0 (IM prime) and 21 (IN boost), female BALB/c mice (n=3 for PBS and n=6 for other PAL-NP groups) were immunized with adjuvanated PAL-NP vaccine formulation with S1 spike protein (see Materials/Methods and Table 1 for doses). Mice were euthanized, and lungs were collected on Day 35 (one-week post-boost). (**A**) Percentage of CD138$^{+}$ASC population. (**B**) Percentage of IgA$^{+}$ASC population. (**C**) Percentage of IgG$^{+}$ASC population. (**D**) Percentage of IgA$^{+}$B$_{RM}$ cell population. (**E**) Percentage of IgG$^{+}$B$_{RM}$ cell population. (**F**) Percentage of GL7$^{+}$ GC B cell population. (**G**) Percentage of IgM$^{+}$ Memory B cell population. Error bars represent the SEM. Statistical significance was calculated with One-Way ANOVA and Tukey post-hoc test. *$p \leq 0.05$, **$p \leq 0.01$, ***$p \leq 0.001$, ****$p \leq 0.0001$ for all graphs.

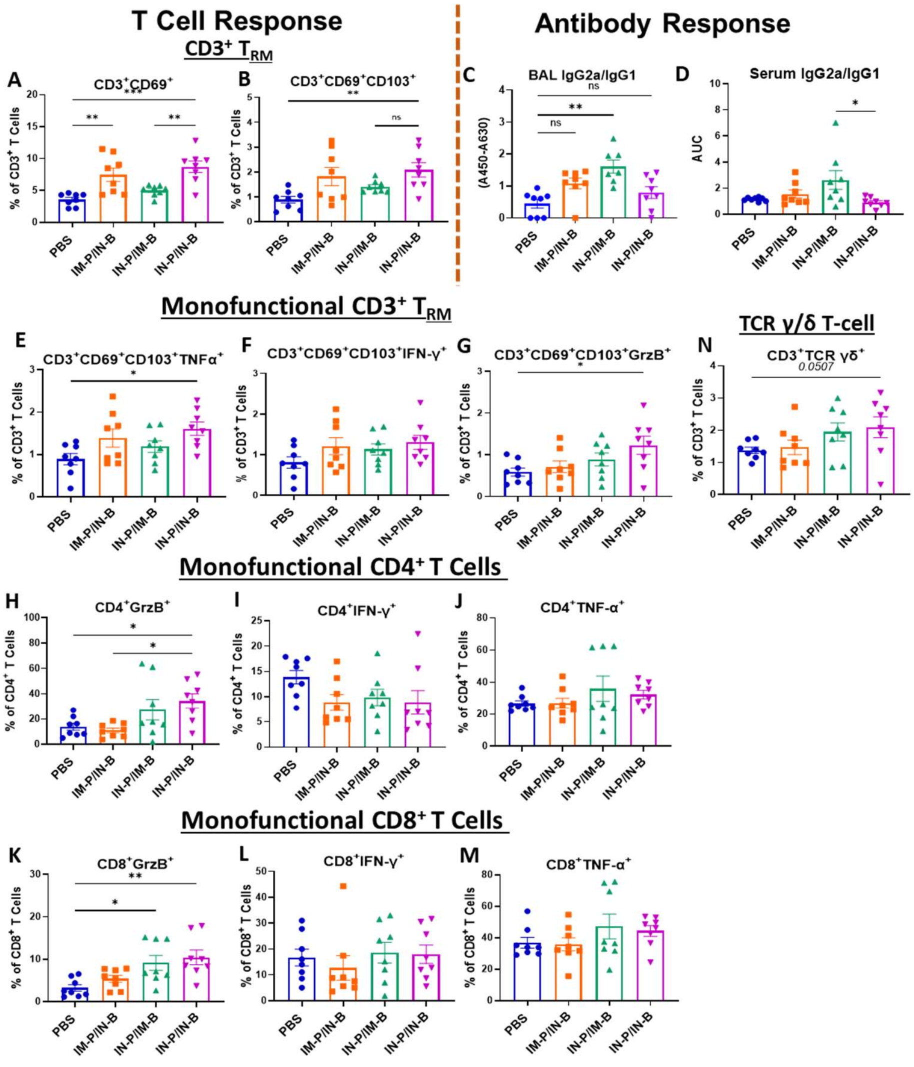


**Figure S5. PUUC+CpG PAL-NP protein subunit vaccine formulation with S1 spike protein, elicits robust SARS-CoV-2 lung-specific T cell immune response with IN-Prime/IN-Boost strategy.** Female BALB/c mice were immunized with PUUC+CpG PAL-NP vaccine formulation with S1 spike protein (see Materials/Methods and Table 1 for doses). Female BALB/c mice (n=8 for all groups) were immunized with three prime-boost strategies: IM-Prime/IN-Boost, IN-Prime/IM-Boost, and IN-Prime/IN-Boost. Mice were euthanized, and lungs were collected on Day 35. Lung cells were restimulated with spike peptide for 6h. (**A**) Percentage of $CD3^+CD69^+$ cell population and, (**B**) percentage of $CD3^+CD69^+CD103^+$ ($CD3^+$ $T_{RM}$) cell population. (**C**) The calculated value of BAL IgG2a/IgG1. (**D**) Calculated value of BAL IgG2a/IgG1. (**E** to **G**) Percentages of monofunctional $CD3^+$ $T_{RM}$ cells expressing TNFα, IFNγ, and GrzB. (**H** to **J**) Percentages of monofunctional $CD4^+$ T cells expressing TNFα , IFNγ, and GrzB. (**K** to **M**) Percentages of monofunctional $CD8^+$ T cells expressing TNFα, IFNγ, and GrzB. (**N**) Percentages of $CD3^+$ TCR γδ cells. (**O**) Percentages of polyfunctional $CD8^+$ $T_{RM}$ cells co-expressing TNF-α and IFN-γ. (**P**) Percentages of polyfunctional $CD8^+$ $T_{RM}$ cells co-expressing TNF-α and IFN-γ. Error bars represent the SEM. Statistical significance was calculated with One-Way ANOVA and Tukey post-hoc test. $*p \leq 0.05$, $**p \leq 0.01$, $***p \leq 0.001$, $****p \leq 0.0001$ for all graphs. Ns represent the non-significant values.

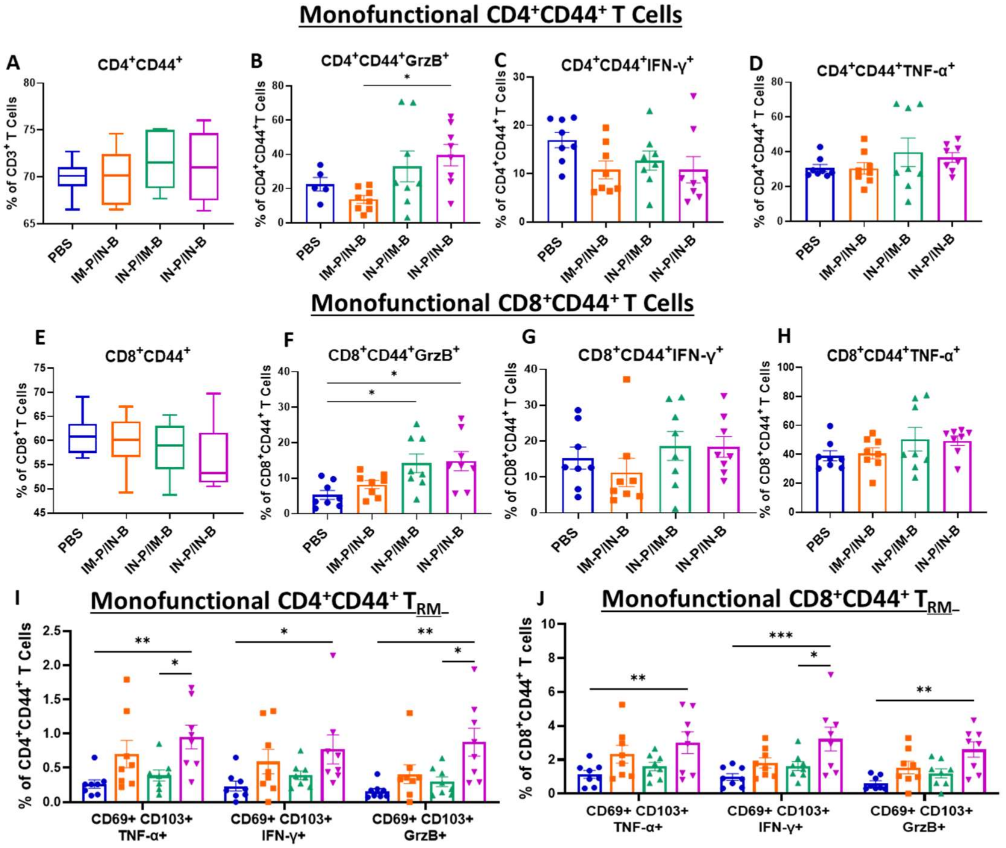


**Figure S6. PUUC+CpG PAL-NP protein subunit vaccine formulation with S1 spike protein, elicits robust SARS-CoV-2 T cell immune responses with IN-Prime/IN-Boost route.** Female BALB/c mice were immunized with PUUC+CpG PAL-NP vaccine formulation with S1 spike protein (see Materials/Methods and Table 1 for doses). Female BALB/c mice (n=8 for all groups) were immunized with three prime-boost strategies: IM-Prime/IN-Boost, IN-Prime/IM-Boost, and IN-Prime/IN-Boost. Mice were euthanized, and lungs were collected on Day 35. Lung cells were restimulated with spike peptide for 6 h. (**A**) Percentages of $CD4^+CD44^+$ cell population. (**B** to **D**) Percentages of monofunctional $CD4^+CD44^+$ cells expressing GrzB, IFN-γ, and TNFα. (**E**) Percentages of monofunctional cells expressing $CD8^+CD44^+$. (**F** to **H**) Percentages of monofunctional $CD8^+CD44^+$ T cells expressing GrzB, IFN-γ, and TNFα. (**I**) Percentages of monofunctional $CD4^+CD44^+$ $T_{RM}$ cells expressing TNFα, IFN-γ, and GrzB. (**J**) Percentages of monofunctional $CD8^+CD44^+$ $T_{RM}$ cells expressing TNFα, IFN-γ, and GrzB. Error bars represent the SEM. Statistical significance T cell frequencies were calculated with One-Way ANOVA and Tukey post-hoc test. $*p \leq 0.05$, $**p \leq 0.01$, $***p \leq 0.001$, $****p \leq 0.0001$ for all graphs.

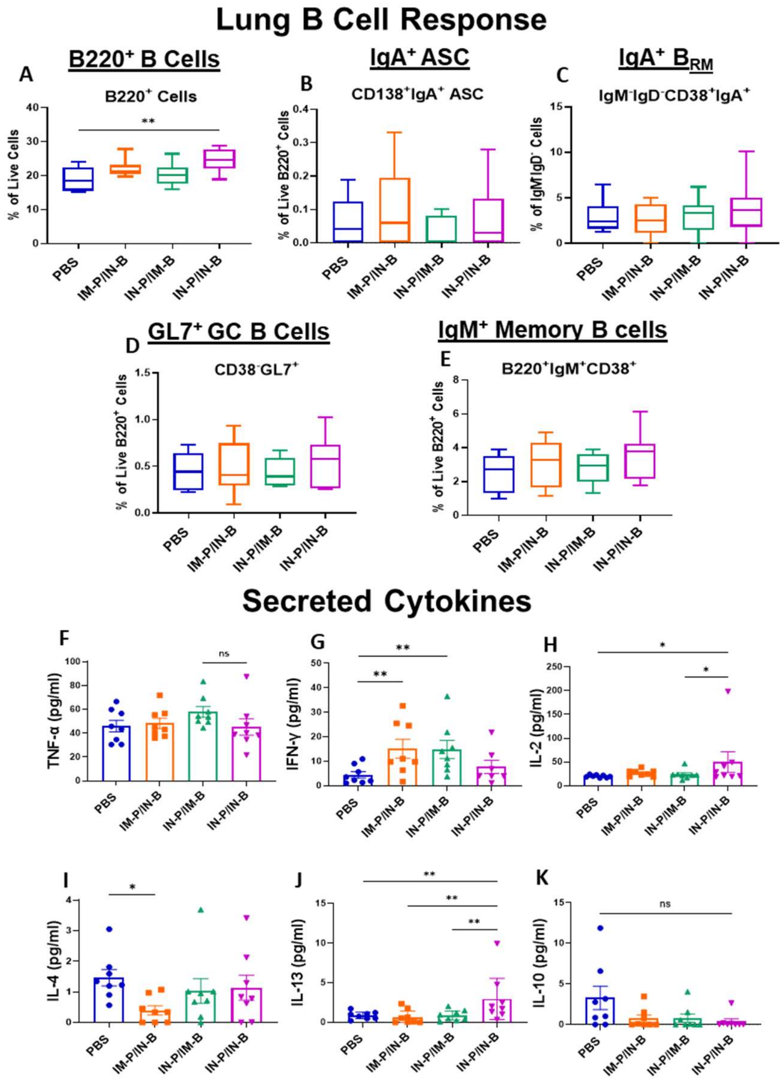


**Figure S7. Lung-specific B cell and T cell (secreted cytokine) responses, when PUUC+CpG PAL-NP protein subunit vaccine formulation and mixed with S1 spike protein, delivered with three different prime-boost routes.** Female BALB/c mice were immunized with PUUC+CpG PAL-NP vaccine formulation with S1 spike protein (see Materials/Methods and Table 1 for doses). Female BALB/c mice (n=8 for all groups) were immunized with three prime-boost strategies: IM-Prime/IN-Boost, IN-Prime/IM-Boost, and IN-Prime/IN-Boost. On days 0 (prime) and 21 (boost), mice were euthanized, and lungs were collected on Day 35. Quantification of B cell response (**A**) Percentage of $B220^+$ B cell population. **(B)** Percentage of $IgA^+$ASC cell population. (**C**) Percentage of $IgA^+$ $B_{RM}$ cell population. (**D**) Percentage of $GL7^+$ GC B cell population. (**E**) Percentage of $IgM^+$ Memory B cell population. Lung cells were restimulated with spike peptide for 6h. (**F** to **K**) Cytokine concentration in supernatants from restimulated lung cells: TNFα, IFN-γ, IL-2, IL-4, IL-13, and IL-10. Error bars represent the SEM. Statistical significance T cell frequencies was calculated with One-Way ANOVA and Tukey post-hoc test. Statistical significance for cytokine concentrations was calculated with one-Way ANOVA and Tukey post-hoc test *$p \leq 0.05$, **$p \leq 0.01$, ***$p \leq 0.001$, ****$p \leq 0.0001$ for all graphs. ns represents the non-significant values.

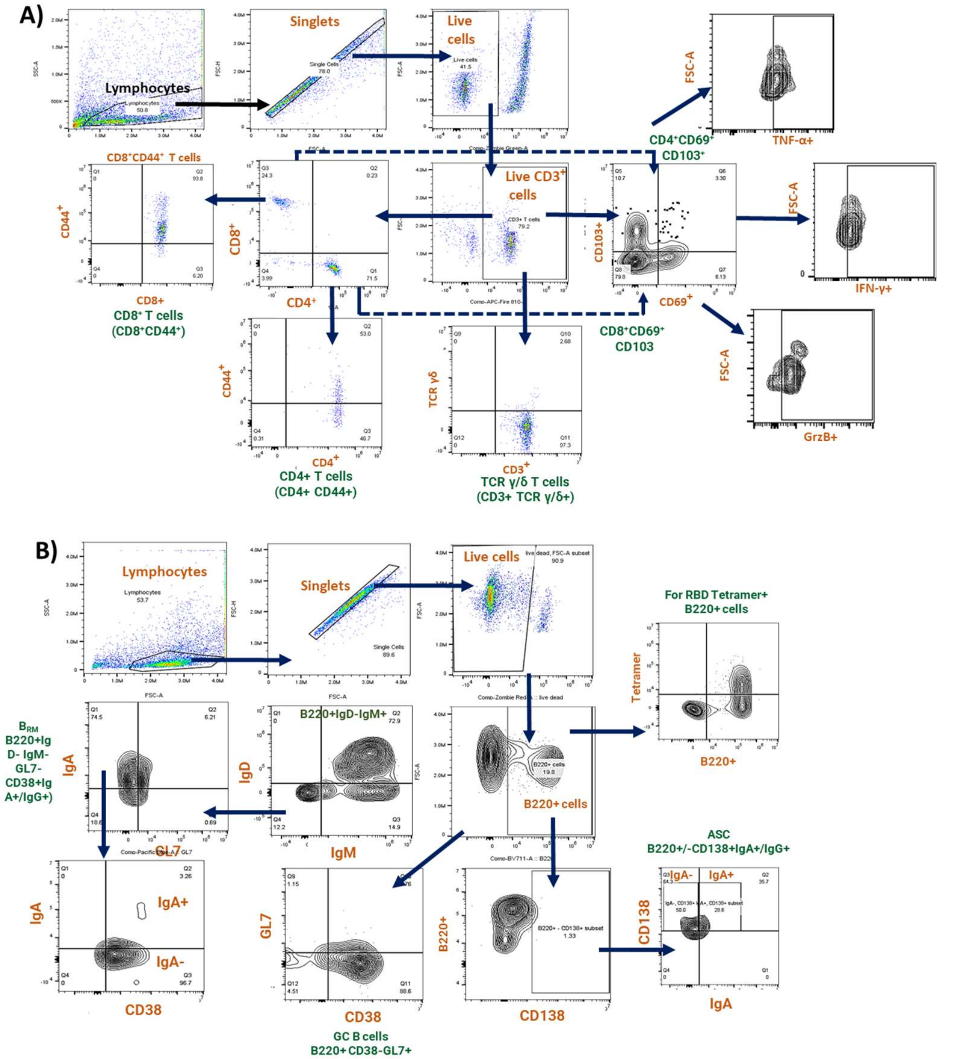


**Figure S8. Gating strategies for analysis of adaptive immune responses in the lungs.** (**A**) Gating strategies to identify CD4$^{+}$ and CD8$^{+}$ T cells and gating strategies to identify cytokine producing CD4$^{+}$ and CD8$^{+}$ T cells. (**B**) Gating strategies to identify antigen-specific and polyclonal B cells.

### NMR Spectrum of Polymers

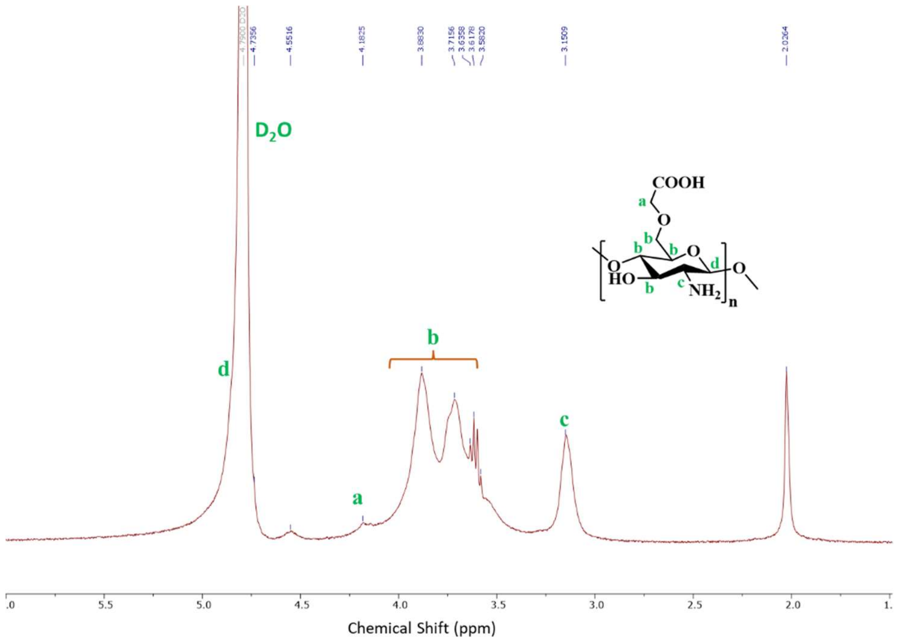


**Figure S9:** 400 MHz $^{1}$H NMR spectrum of the carboxylated chitosan (OCMC) in $D_2O$ with 1% DCl.

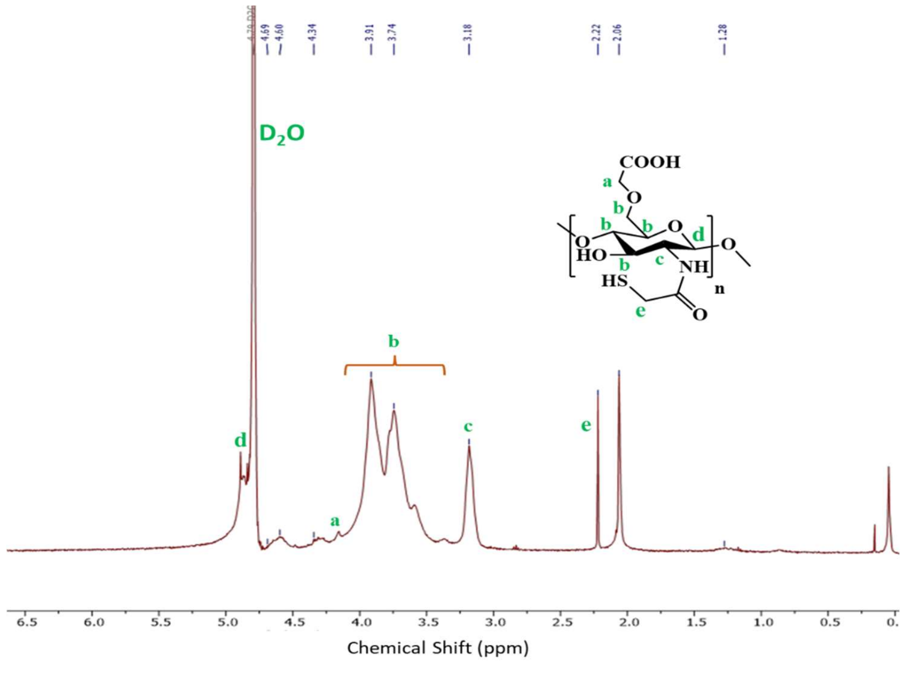


**Figure S10:** 400 MHz $^{1}$H NMR spectrum of the thiolated OCMC in $D_2O$ with 1% DCl.

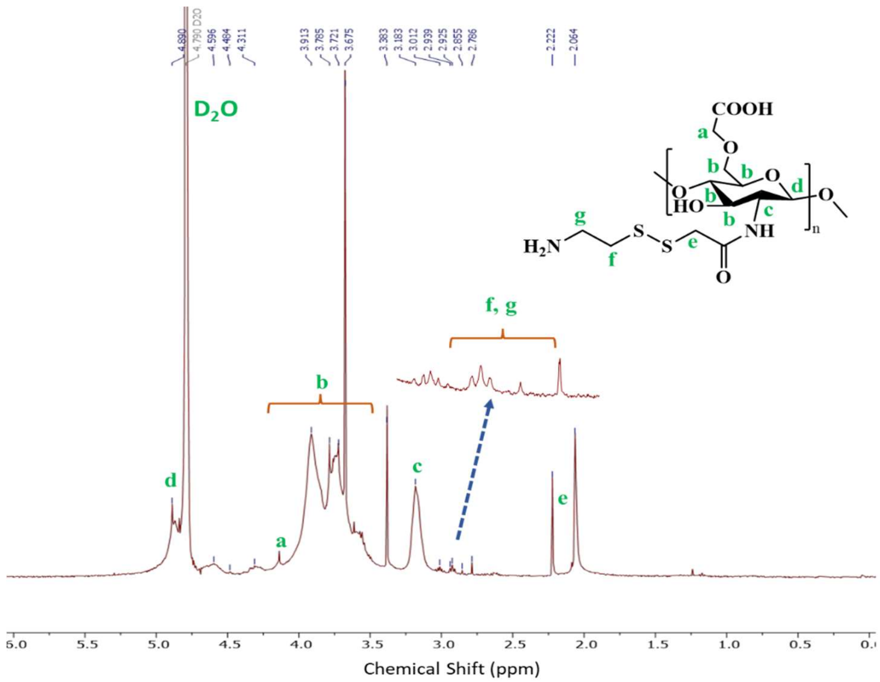


**Figure S11:** 400 MHz $^1$H NMR spectrum of the OCMC-S-S-Cys in $D_2O$ with 1% DCl.

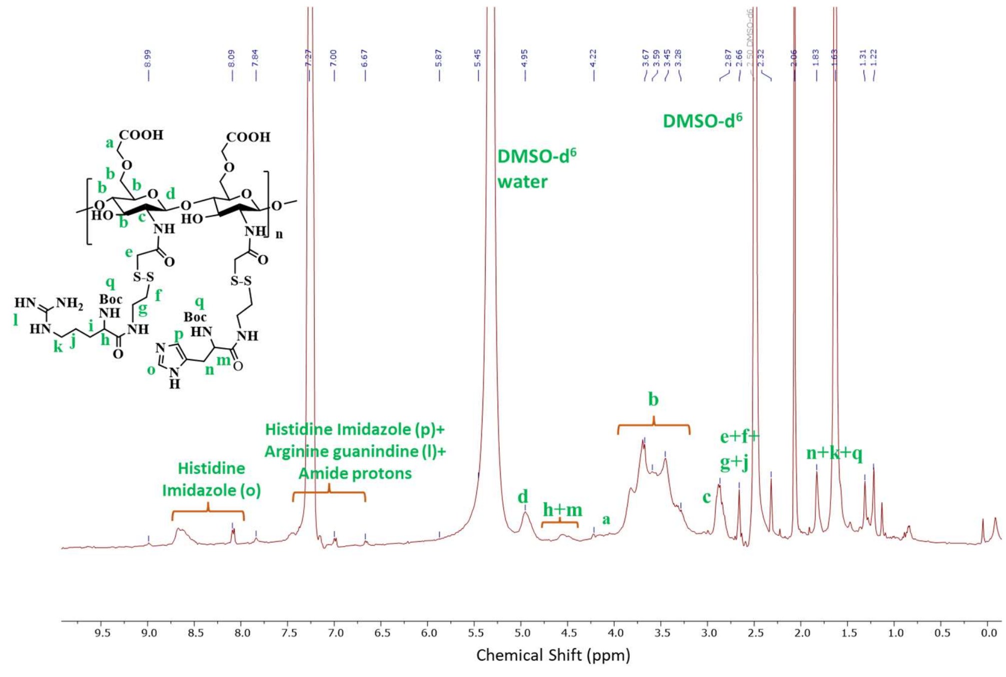


**Figure S12:** 400 MHz $^1$H NMR spectrum of the OCMC-S-S-(A/H) in DMSO-d$^6$..

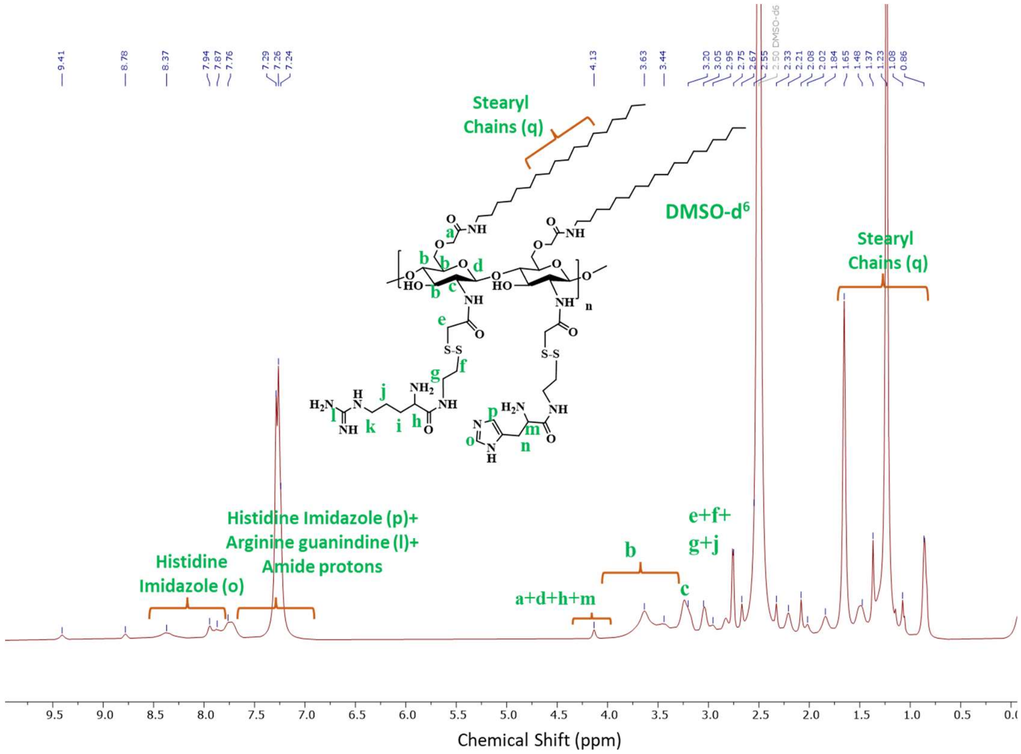


**Figure S13:** 400 MHz $^1$H NMR spectrum of the OCMC-S-S-(A/H)-SA in DMSO-d$^6$.
